# Modeling the Evolution of Rates of Continuous Trait Evolution

**DOI:** 10.1101/2022.03.18.484930

**Authors:** B. S. Martin, G. S. Bradburd, L. J. Harmon, M. G. Weber

## Abstract

Rates of phenotypic evolution vary markedly across the tree of life, from the accelerated evolution apparent in adaptive radiations to the remarkable evolutionary stasis exhibited by so-called “living fossils”. Such rate variation has important consequences for large-scale evolutionary dynamics, generating vast disparities in phenotypic diversity across space, time, and taxa. Despite this, most methods for estimating trait evolution rates assume rates vary deterministically with respect to some variable of interest or change infrequently during a clade’s history. These assumptions may cause underfitting of trait evolution models and mislead hypothesis testing. Here, we develop a new trait evolution model that allows rates to vary gradually and stochastically across a clade. Further, we extend this model to accommodate generally decreasing or increasing rates over time, allowing for flexible modeling of “early/late bursts” of trait evolution. We implement a Bayesian method, termed “evolving rates” (*evorates* for short), to efficiently fit this model to comparative data. Through simulation, we demonstrate that *evorates* can reliably infer both how and in which lineages trait evolution rates varied during a clade’s history. We apply this method to body size evolution in cetaceans, recovering substantial support for an overall slowdown in body size evolution over time with recent bursts among some oceanic dolphins and relative stasis among beaked whales of the genus *Mesoplodon*. These results unify and expand on previous research, demonstrating the empirical utility of *evorates*.

---

The rates at which traits evolve is markedly heterogeneous across the tree of life, as evidenced by the uneven distribution of phenotypic diversity across space, time, and taxa (e.g., Simpson, 1944; Reaney et al., 2020; Brusatte et al., 2012; Chartier et al., 2021). While understanding the drivers of such patterns can provide critical insights into macroevolutionary processes, general consensus on what factors are most important in accelerating and decelerating trait evolution remain elusive (Chira et al., 2018). There is a vast, interconnected web of factors hypothesized to affect trait evolution rates, typically divided into extrinsic and intrinsic components. Extrinsic factors relate to the environment of an evolving lineage, commonly including aspects of biogeography like climate or habitat (e.g., Clavel and Morlon, 2017; Mihalitsis and Bellwood, 2019), as well as interactions with other species (e.g., Slater, 2015; Borstein et al., 2019; Drury et al., 2021). Intrinsic factors instead involve properties of the evolving lineage itself, including life history attributes such as behavior or developmental traits (e.g., Muñoz and Bodensteiner, 2019; Fabre et al., 2020) and genetic features like trait heritability and effective population size (e.g., Arnold et al., 2008; Villar et al., 2014). The effects of all these variables are interrelated and depend on the particular traits being studied, further complicating matters (Cooper and Purvis, 2009; Muñoz et al., 2018; see also Donoghue and Sanderson, 2015).

Unfortunately, the evolutionary histories of many factors hypothesized to affect trait evolution rates are largely unobserved. Thus, methods testing for associations between rates and variables of interest must first estimate the history of the explanatory variables themselves (but see Hansen et al., 2021). This limits researchers to considering only a few, relatively simple hypotheses (Revell, 2013; Caetano and Harmon, 2019), causing most trait evolution models to underfit observed data (see also Pennell et al., 2015). This underfitting generally oversimplifies inferred rate variation patterns and artificially increases statistical support for complex models which may imply spurious links between trait evolution rates and explanatory variables (May and Moore, 2020; see also Rabosky and Goldberg, 2015; Beaulieu and O’Meara, 2016). Thus, these “hypothesis-driven” approaches to modeling trait evolution should be integrated with “data-driven” approaches that agnostically model variation in trait evolution rates based on observed trait data alone. Such approaches can account for rate variation unrelated to some focal hypothesis, or even be used to generate novel hypotheses regarding what factors may have driven inferred rate variation patterns (Uyeda et al., 2018; May and Moore, 2020; see also Beaulieu and O’Meara, 2016).

Several data-driven methods for inferring trait evolution rates are already available and widely used (Eastman et al., 2011; Thomas and Freckleton, 2012; Rabosky et al., 2014; Pagel et al., 2022), but such methods generally work by splitting phylogenetic trees into subtrees and assigning a unique rate to each subtree (sometimes termed “macroevolutionary regimes”). These models implicitly assume trait evolution rates stay constant over long periods of time with sudden shifts in particular lineages. This mode of rate variation would be expected if rates are primarily influenced by only a few, discretely varying factors of large effect. However, this assumption could be problematic given the sheer number of factors hypothesized to affect trait evolution rates, as well as the fact that many of these factors vary continuously (Cooper and Purvis, 2009). If rates are instead affected by many factors, mostly with subtle effects, we would expect trait evolution rates to constantly shift in small increments over time within a given lineage, resulting in gradually changing rates over time and phylogenies. In other words, rates themselves would “evolve” and be similar, but not identical, among closely-related lineages (i.e., phylogenetic autocorrelation; see Sakamoto and Venditti, 2018). By assuming that rates change infrequently, current data-driven methods likely oversimplify rate variation patterns, collapsing heterogeneous evolutionary processes into homogeneous regimes (but see May and Moore, 2020; Fisher et al., 2021). To this end, Revell (2021) recently developed a data-driven method that models trait evolution as gradually changing, but this method is limited in requiring *a priori* specification of how much trait evolution rates vary across the phylogeny. Further, the method offers no way to rigorously test whether lineages exhibit significantly different rates (Revell, 2021).

Notably, some hypothesis-driven methods model trait evolution rates as gradually changing over time. However, such models most commonly assume that rates only follow a simple trend of exponential decrease or increase over time (Blomberg et al., 2003; but see Clavel and Morlon, 2017; Slater et al., 2017). In this context, declining trait evolution rates, or “early bursts” (EB), are often invoked as signatures of adaptive radiation (Harmon et al., 2010), while increasing trait evolution rates, or “late bursts” (LB), are sometimes linked to processes like character displacement (Weber et al., 2016; Skeels and Cardillo, 2019). Unfortunately, current methods lack statistical power to detect decreasing trends in rates when just a few lineages deviate from an overall EB pattern (Slater and Pennell, 2014). Essentially, by assuming a perfect correspondence between time and rates across all lineages, inference under these methods is misled by subclades exhibiting anomalously low or high trait evolution rates. New methods that explicitly model such “residual” rate variation may more accurately detect general trends in trait evolution rates by accounting for these anomalous lineages/subclades.

Here we develop a new, data-driven method, called “evolving rates” (*evorates* for short), that models trait evolution rates as gradually changing over time, ultimately resulting in stochastic, continuously distributed rates that are more similar among closely-related lineages. We take advantage of recent developments in Bayesian inference and develop new strategies for efficiently estimating autocorrelated rates on phylogenetic trees while dealing with uncertain trait values, resulting in relatively fast, reliable inference. *Evorates* is flexible and intuitive, allowing researchers to infer both how and where rates vary on a phylogeny. Through simulation, we demonstrate that *evorates* recovers accurate parameter estimates on ultrametric phylogenies spanning a range of sizes and that it is more sensitive and robust in detecting trends in trait evolution rates than conventional EB/LB models. We also use *evorates* to model body size evolution among extant whales and dolphins (order cetacea) and find evidence for declining rates of body size evolution and moderate rate heterogeneity in this clade, unifying and expanding on previous results (Slater et al., 2010; Slater and Pennell, 2014; Sander et al., 2021).

## Materials and Methods

*Evorates* uses comparative data on a univariate continuous trait to infer how trait evolution rates change over time as well as which lineages in a phylogeny exhibit anomalous rates. Here, comparative data refers to a fixed, rooted phylogeny with branch lengths proportional to time and trait values associated with its tips. The method is designed to work with raw trait measurements; both missing data and multiple trait values per tip are allowed (i.e., tips with 0 and *>* 1 observations, respectively). However, users may also use estimated mean trait measurements by assigning a given tip a single value with associated standard error, essentially specifying a normal prior for the trait value at that tip. While we focus on ultrametric phylogenies in the current paper, we note that the method also works with non-ultrametric phylogenies as well; such priors might be based on incomplete fossil data and even assigned to internal nodes of the phylogeny (see Slater et al., 2012). Conditional on these trait data, *evorates* uses Bayesian inference to estimate two key parameters governing the process of rate change: rate variance, controlling how quickly rates diverge among independently evolving lineages, and a trend, determining whether rates tend to decrease or increase over time. When rate variance is 0, rates do not accumulate random variation over time and are constant across contemporaneous lineages. In this case, trait evolution follows an EB process if the trend is negative, Brownian Motion (BM) if it is 0, or LB if it is positive. The method also infers branchwise rates, which are estimates of average trait evolution rates along each branch in the phylogeny, indicating which lineages exhibit unusually low or high rates.

### The Model

At its core, *evorates* works by extending a typical Brownian Motion (BM) model of univariate trait evolution to include stochastic, incremental changes in trait evolution rates, *σ*^2^. Specifically, *σ*^2^ follows a process approximating geometric BM (GBM) with a constant rate, meaning that ln(*σ*^2^) follows a homogeneous BM-like process. GBM is a natural process to describe “rate evolution” since it ensures rates stay positive and implies rates vary on a multiplicative, as opposed to additive, scale (Limpert et al., 2001; Gingerich, 2009). To render inference under this model tractable, we treat it as a hierarchical model with a trait evolution process dependent on the unknown–but estimable–branchwise rates, which are themselves dependent on a rate evolution process controlled by the estimated rate variance and trend parameters. The overall posterior probability of the model can be summarized as:

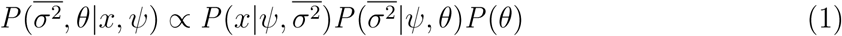

Where *ψ* is a phylogeny with *e* branches and *n* tips, 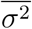 is an *e*-length vector of branchwise rates, *x* is an *n*-length vector of trait values for each tip, and *θ* is a vector of parameters governing the rate evolution process. Cases with missing data and multiple trait values per tips are covered in a later section. In our notation, time is 0 at the root of the phylogeny and increases towards the tips. 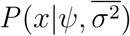 is the likelihood of *x* given the trait evolution process, 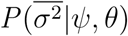 is the probability of branchwise rates given the rate evolution process, and *P* (*θ*) is the prior probability of the rate evolution process parameters. We explicitly estimate and condition likelihood calculations on branchwise rates (a type of “data augmentation”; see May and Moore, 2020) since the likelihood of the trait data while marginalizing over branchwise rates (i.e., *P* (*x*|*ψ, θ*)) does not follow a known probability distribution and would require complex, numerical approximations to compute. On the other hand, 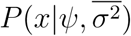 follows a straight-forward multivariate normal density:

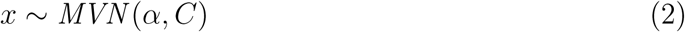

where *α* is a vector of the trait value at the root of the phylogeny repeated *n* times and *C* is an *n × n* matrix. The entries of *C* are given by:

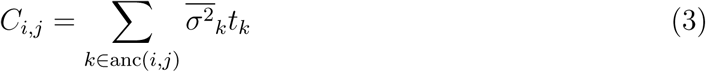

where *t* is an *e*-length vector of branch lengths, *i* and *j* are indices denoting specific tips, *k* is an index denoting a particular branch, and anc(*i, j*) is a function that returns all ancestral branches shared by *i* and *j*. Note that when branchwise rates are constant across the tree, *C*_*i,j*_ is proportional to the elapsed time between the root of the phylogeny and the most recent common ancestor of *i* and *j*. Branchwise rates can be thought of as “squashing” and “stretching” the branch lengths of a phylogeny, such that certain lineages have evolved for effectively shorter or longer amounts time, respectively.

Unfortunately, there is no general solution for calculating 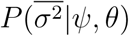 under a true GBM process (Lepage et al., 2007), so we instead use a multivariate log-normal approximation (e.g., Dufresne, 2004; Welch and Waxman, 2008) of the distribution of branchwise rates and calculate probabilities under this approximation. Briefly, this approximation decomposes branchwise rates into their expected values, *β*, determined solely by the trend parameter, and a “noise” component, *γ*, sampled from a multivariate normal distribution controlled by the rate variance parameter:

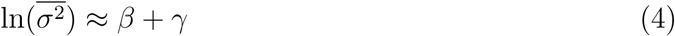

Here, the noise component is approximate since it follows the distribution of geometric, rather than arithmetic, averages of trait evolution rates along each branch assuming there is no trend (i.e., 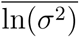) rather than 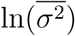; see online appendix for further details). The entries of *β* are given by:

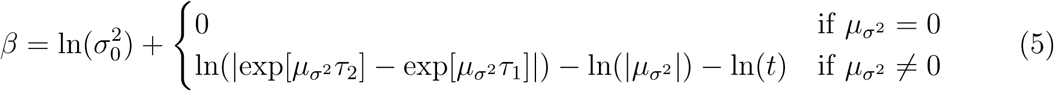

where 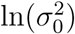 is the estimated rate at the root of the phylogeny, 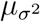 is the trend parameter, *t* is an *e*-length vector of branch lengths, and *τ*_1_ and *τ*_2_ are *e*-length vectors of the start and end times of each branch in the phylogeny (Blomberg et al., 2003). The entries of *γ* are given by:

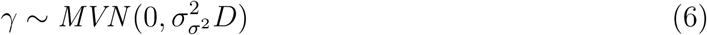

where 0 is a vector of 0s repeated *e* times, 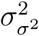 is the rate variance parameter, and *D* is an *e × e* matrix. The entries of *D* are given by:

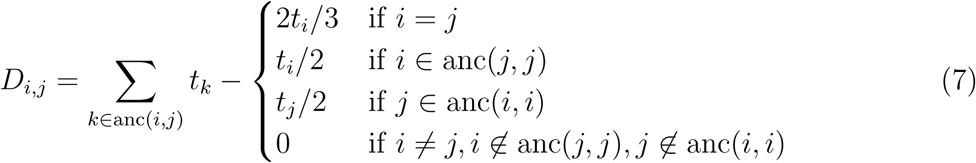

where *i, j*, and *k* are all indices denoting branches and anc(*i, j*) is a function that returns all ancestral branches shared by *i* and *j* (Devreese et al., 2010; see online appendix for further details). Overall, this approximation closely matches the distribution of branchwise rates obtained via fine-grained simulations of GBM on phylogenies under plausible parameter values and is negligibly different from other computationally efficient approximations (e.g., Thorne et al., 1998; Lartillot and Poujol, 2011; Revell, 2021; Figs. S3-16; Tables S2-4). We prefer this approximation because it is convenient to work with and directly focuses on estimating branchwise rates rather than rates at the nodes of the phylogeny, which is what other strategies focus on.

Under this approximation, the final expression for the posterior probability is:

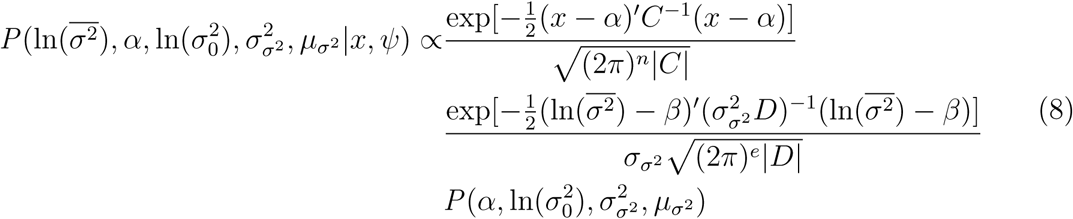

### Model Implementation

*Evorates* estimates the posterior distribution of parameters given a phylogeny and associated trait data via Hamiltonian Monte Carlo (HMC) using the probablistic programming language Stan, interfaced through R (Carpenter et al., 2017; Stan Development Team, 2020, 2021). To optimize sampling efficiency and avoid numerical issues, *evorates* estimates branchwise rates with an uncentered parameterization (Betancourt and Girolami, 2013) and marginalizes over unobserved trait values at the root and tips of the tree (Freckleton, 2012; Hassler et al., 2020). Under an uncentered parameterization, the HMC algorithm does not directly estimate branchwise rates, but instead estimates the distribution of *e* independent standard normal random variables, *z*, which are transformed to follow the distribution of branchwise rates:

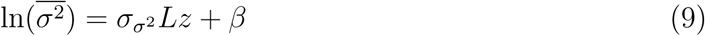

where *L* is lower triangular Cholesky factorization of *D* (i.e., *D* = *LL*^*′!*^; see Eqn. 7). This parameterization is particularly efficient since it avoids having to repeatedly manipulate *D* to calculate 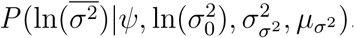.

*Evorates* also uses Felsenstein’s pruning algorithm for quantitative traits to marginalize over the trait value at the root of the phylogeny and avoid repeatedly inverting *C* when calculating 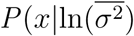 (Felsenstein, 1973; Freckleton, 2012; Caetano and Harmon, 2019). To simplify the pruning algorithm implementation, any multifurcations in the phylogeny are converted to series of bifurcations by adding additional “psuedo-branches” of length 0. Importantly, this procedure does not alter the resulting likelihood calculations (Felsenstein, 2008). While branchwise rates along these pseudo-branches are identifiable based on the rate evolution process, our implementation does not estimate them since they do not affect the likelihood of observed trait data and are not particularly interesting from an empirical standpoint. We also modify Felsenstein’s pruning algorithm to marginalize over unobserved trait values at the tips, including cases involving standard error, missing data, and multiple observations per tip.

### Accommodating Missing Data and Multiple Observations

Incorporating uncertainty in observed trait values in comparative studies is especially important for methods that model trait evolution rate variation, since measurement error can inflate estimates of evolutionary rates, particularly in young clades (Felsenstein, 2008). To prevent such biases, *evorates* generally treats the true (mean) trait values at the tips, *x*, as an unknown parameter. We marginalize over *x* given raw trait measurements, *y* (potentially including 0 or *>* 1 observations for some tips), and “tip error” variances for each tip, 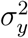. While we use the term “raw” trait measurement for clarity, the data provided for certain tips could instead be the average of multiple trait measurements or even the mean of a normal prior on the trait value. Entries of 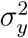 for such tips may be fixed to associated standard error or prior variances. All other entries of 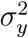 are treated as unfixed, free parameters. To render the model more tractable, we assume tip error variance is constant across all tips with unfixed variance.

To marginalize over the true trait values, we modify the initialization of Felsenstein’s pruning algorithm (Felsenstein, 1973). Prior to pruning, we assign each tip the expectation and variance of the true trait value given the raw trait measurements. We then calculate each tip’s partial likelihood from contrasts between its associated raw trait measurements given its error variance, 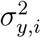. Assuming the raw trait measurements are independently sampled from a normal distribution with variance 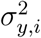, the true trait value’s expectation is simply the mean of the raw trait measurements, 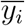, and its variance is given by 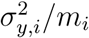, where *m*_*i*_ is the number of raw trait measurements (Felsenstein, 2008). Note that if there are no trait measurements for a particular tip (i.e., *m*_*i*_ = 0), the expectation of that tip’s true trait value is undefined with infinite variance (Hassler et al., 2020).

Since there are no contrasts for tips with one or fewer raw trait measurements, the partial likelihood associated with these tips is 1. Otherwise, we can derive a general formula for the partial likelihood by considering each tip as a small subtree and applying Felsenstein’s pruning algorithm. Specifically, each tip is treated as a star phylogeny consisting of *m*_*i*_ “sub-tips” of length 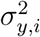, with trait values *y*_*i*_ (Felsenstein, 1973, 2008):

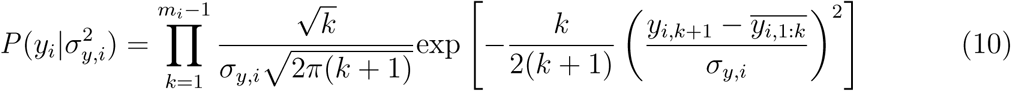

where *i* denotes a particular tip, *y*_*i*_ is a vector of *m*_*i*_ raw trait measurements for tip 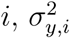 is the tip error variance for tip *i*, and 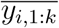 is the mean of measurements 1 through *k* in the vector *y*_*i*_.

After initializing all tips in the phylogeny, Felsenstein’s pruning algorithm can be applied normally, iterating over the internal nodes from the tips towards the root (e.g., Felsenstein, 1973, Freckleton, 2012, Caetano and Harmon, 2019). The presence of missing data, however, will cause some calculations to involve nodes with undefined expected trait values and infinite variance. Note that these “data-deficient” nodes do not contribute information to the expectation and variance of the trait value at their ancestral nodes. Thus, if both nodes descending from some focal node are data deficient, the focal node will also be data deficient, with undefined expectation and infinite variance. Otherwise, if only one descendant node is data deficient, the expectation and variance of the trait value at the focal node is solely determined by the descendant node that is *not* data deficient. Let the descendant, non-data deficient node have expected trait value and variance 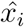 and 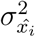, respectively, and be connected to the focal node by a branch of length *t*_*i*_ with branchwise rate 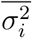. The focal node’s expected trait value and variance will be 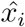 and 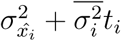, respectively. Whether one or both descendant nodes are data deficient, there is no contrast associated with the focal node and the corresponding partial likelihood is 1.

In the case of univariate traits, the benefit of including missing data is that ancestral state reconstruction algorithms can be used to estimate the trait values of tips with missing data (Goolsby, 2017; Hassler et al., 2020). Tips with missing data have no effect on the likelihood or parameter estimation, other than perhaps improving how well the rate evolution process approximates GBM in limited cases.

### Priors

In HMC-based Bayesian inference, uninformative priors tend to result in fat-tailed posterior distributions and produce numerical issues as the sampler explores unrealistic regions of parameter space (Stan Development Team, 2021). We thus chose weakly informative priors that preserve the overall location and precision of posterior distributions while preventing these sampling problems. Let 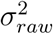 be the variance of the provided trait data and *T* the overall height of the phylogeny. A normal prior with mean 0 and standard deviation 10*/T* is placed on the trend parameter 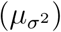. A log-normal prior with location 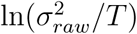 and scale 10 is placed on the rate at the root 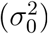. Half-cauchy priors with scales 5*/T* and 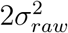 are placed on the rate and unfixed tip error variances (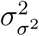 and 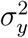), respectively. Note that most of these parameters are sampled on a log scale, but affect trait evolution dynamics on an exponential scale. Given an ultrametric phylogeny, a trend of 10*/T* corresponds to a *∼*20,000-fold increase in trait evolution rates from the root to tips, and data simulated with a rate variance of 5*/T* typically accumulate a *∼*1,000 to 10,000-fold rate difference between the most slowly and quickly evolving tips.

### Significance Testing

We agree with other macroevolutionary biologists advocating for greater focus on interpreting parameter estimates and effect sizes inferred by comparative models (e.g., Beaulieu and O’Meara, 2016). Nonetheless, assessing statistical support for particular hypotheses remains important for biologically interpreting fitted models. In the context of *evorates*, we focus on two main hypotheses: 1) that significant rate heterogeneity, independent of any trend, occurred over the history of a clade 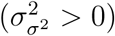, and 2) rates generally declined or increased over time (i.e., 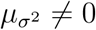). Both hypotheses could be tested by fitting additional models with constrained rate variance and/or trend parameters and comparing among unconstrained and constrained models using Bayes factors. However, Bayes factor estimation requires additional, time-consuming computation. Thus, we developed alternative hypothesis-testing approaches that only require the posterior samples of a fitted, unconstrained model.

We use the posterior probability that 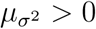 to test for overall trends in rates. If the posterior probability is 0.025 or less, we can conclude that there is substantial evidence that rates declined over time, and vice versa if the posterior probability is 0.975 or above. This corresponds to a two-tailed test with a critical value of 0.05. For rate variance, we instead use Savage-Dickey (SD) ratios since rate variance is bounded at 0 and the posterior probability that 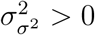 will always be 1. SD ratios are ratios of the posterior to prior probability density at a particular parameter value corresponding to a null hypothesis. If this ratio is sufficiently less than 1, the data have “pulled” prior probability mass away from the null hypothesis, suggesting that the null hypothesis is likely incorrect. In general, a ratio of 1*/*3 or less is considered substantial evidence against the null hypothesis (Kass and Raftery, 1995). We use log spline density estimation implemented in the R package *logspline* (version 2.1.16) to estimate the posterior probability density at 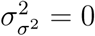 (Stone et al., 1997; Wagenmakers et al., 2010).

Researchers may also wish to identify lineages evolving at anomalous rates. The most straight-forward method to do so is to calculate the posterior probability that branchwise rates are greater than some “background rate”, analogous to the approach for trends. In this paper, we define the background trait evolution rate as the geometric mean of branchwise rates, weighted by their relative branch lengths. Rates are generally distributed with long right tails (Gingerich, 2009), particularly under our model whereby rate evolution follows a GBM-like process. Geometric means are less sensitive than arithmetic means to extremely high, outlier rates associated with these long tails, and are thus better-suited for rate comparisons. Note that we can also calculate background rates for subsets of branches in a phylogeny, such that trait evolution rates for subclades can be estimated and compared.

In the presence of a strong trend, lineages with anomalous rates will occur at an ultrametric phylogeny’s root and tips, rendering anomalous rate detection redundant with trend estimation. Thus, we define a helpful branchwise rate transformation, called “detrending”, which further facilitates interpretation of *evorates* results. Specifically, branchwise rates are detrended prior to calculating background rates and posterior probabilities by subtracting *β* from branchwise rates on the natural log scale (see Eqn. 5). These detrended rates yield a new set of transformed parameters, branchwise rate deviations, 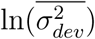, defined as the difference between detrended branchwise rates and the background detrended rate on the natural log scale. When the posterior probability 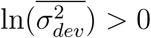 for a given branch is less than 0.025 or greater than 0.975, we can conclude that trait evolution is anomalously slow or fast along that branch, respectively, given the overall trend in rates through time. While we focus on comparing detrended branchwise and background rates based on geometric means in the current paper, we note that *evorates* can also compare untransformed branchwise and background rates based on either geometric or arithmetic means per user specifications.

### Simulation Study

To test the performance and accuracy of *evorates*, we applied it to continuous trait data simulated under the model of inference. We simulated data under all combinations of no, low, and high rate variance 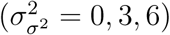 and decreasing, constant, and increasing trends 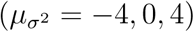, for a total of 9 trait evolution scenarios. We picked these values to simulate data that appeared empirically plausible and represented a range of different trait evolution dynamics. Note that when rate variance is 0, the resulting simulations evolve under EB, BM, or LB models of trait evolution depending on the trend parameter. We simulated traits evolving along ultrametric, pure-birth phylogenies with 50, 100, and 200 tips generated using the R package *phytools* (version 1.0-1; Revell, 2012) to assess the effect of increasing sample size on model performance. We simulated 10 phylogenies and associated trait data for each trait evolution scenario and phylogeny size for a total of 270 simulations. In all cases, phylogenies were rescaled to a total height of 1, ensuring the effect of parameters remained consistent across replicates. All simulations were simulated with a trait and log rate value of 0 at the root. Since we focused on the estimation of branchwise rate, rate variance, and trend parameters, we simulated trait data with only 1 observation per tip and no tip error.

To quantitatively assess the simulation study results, we calculated the breadth, relative accuracy, and coverage of marginal posterior distributions for rate variance and trend parameters. Here, breadth is the width of 95% credible interval, while relative accuracy is absolute difference between the posterior distribution median and true value, divided by the corresponding breadth. Coverage is a binary metric equal to one when the true value falls within the 95% credible interval and zero otherwise. For branchwise rate parameters, we averaged the breadth, relative accuracy, and coverage of all branchwise rate marginal posterior distributions (on the natural log scale) for each model fit. Additionally, we calculated the statistical power and false positive error rate (i.e., type I error rate, hereafter error rate) of *evorates* for detecting significant rate variance and decreasing/increasing trends. Due to the continuous nature of branchwise rates, we assessed power and error rates for detecting anomalous branchwise rates by calculating the proportion of times a branch is detected as exhibiting anomalously slow or rapid trait evolution rates across different values of true branchwise rate deviations.

### Empirical Example

We applied *evorates* to model body size evolution in extant cetaceans using the most recently estimated timetree of both fossil and extant cetaceans (Lloyd and Slater, 2021), pruned to consist of 88 extant species (we excluded one extant species, *Balaenoptera brydei*, due to its uncertain taxonomic status; see Constantine et al., 2018), and associated trait data on log-transformed maximum female body lengths for each species. Most body length data was compiled in a previous comparative study, but we supplemented these data with published measurements for an additional 15 species (Table S1). We chose this example since previous research detected notable signatures of declining body size evolution rates over time in this clade, despite conventional model selection failing to yield support for an EB model of trait evolution. This puzzling result seems primarily due to a few recently-evolved lineages exhibiting unusually rapid shifts in body size (Slater et al., 2010; Slater and Pennell, 2014; see also Sander et al., 2021). While previous work used a mix of simulation and outlier detection techniques to arrive at this conclusion, we predicted that our method would identify these patterns in a more cohesive modeling framework.

### HMC configuration and diagnostics

When fitting models to simulated and the empirical data, we ran 4 HMC chains consisting of 3,000 iterations. After discarding the first 1,500 iterations as warmup and checking for convergence, chains were combined for a total of 6,000 HMC samples for each simulation. We repeated this procedure while constraining the rate variance parameter to 0 to see if our method could detect trends in trait evolution rates with more power than conventional EB/LB models. We set tip error for the simulation study to 0 *a priori* since we do not focus on inference of this parameter here, though we did allow the method to estimate tip error in the empirical example. For each model fit, chains mixed well (greatest 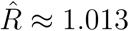) and achieved effective sample sizes of at least 3,000 for every parameter. Divergent transitions, a feature of HMC which can be indicative of sampling problems, were relatively rare, with only six simulation model fits exhibiting 1-3 divergent transitions. Overall, diagnostic tests suggested all HMC chains converged and sampled posterior distributions thoroughly.

## Results

### Performance of Method

Overall, the method exhibited good coverage and relative accuracy for all parameters, though posterior distribution breadth was often quite large, especially for trees with 50 tips (Tables 1-3, Fig. 1). Breadth was highly dependent on trait evolution scenario and tree size. In general, higher values of trends and rate variance were associated with wider posteriors for their respective parameters, such that increasing trends and high rate variance are estimated with the least precision. In some cases, higher trends seemed to increase the breadth of rate variance posteriors and vice versa, but this pattern was weak overall. Despite the shrinking breadth of posteriors with larger tree size, coverage for trend and rate variance parameters across all trait evolution scenarios and tree sizes remained consistent at around the theoretical expectation of 95%.

**Table 1.** Average posterior relative accuracy for rate variance, trend, and branchwise rate parameters for each simulated trait evolution scenario and tree size

| $\sigma_{\sigma^2}^2 =$ | | rate variance | | | trend | | | branchwise rates | | |
| --- | --- | --- | --- | --- | --- | --- | --- | --- | --- | --- |
|  |  | 0 | 3 | 6 | 0 | 3 | 6 | 0 | 3 | 6 |
| 50 species |  |  |  |  |  |  |  |  |  |  |
| $\mu_{\sigma^2} =$ | -4 | 0.17 | 0.23 | 0.11 | 0.19 | 0.16 | 0.22 | 0.13 | 0.21 | 0.21 |
|  | 0 | 0.15 | 0.24 | 0.24 | 0.21 | 0.20 | 0.25 | 0.14 | 0.20 | 0.22 |
|  | 4 | 0.21 | 0.15 | 0.27 | 0.12 | 0.22 | 0.25 | 0.12 | 0.19 | 0.22 |
| 100 species |  |  |  |  |  |  |  |  |  |  |
| $\mu_{\sigma^2} =$ | -4 | 0.18 | 0.11 | 0.22 | 0.17 | 0.18 | 0.23 | 0.12 | 0.19 | 0.22 |
|  | 0 | 0.19 | 0.24 | 0.25 | 0.23 | 0.15 | 0.19 | 0.13 | 0.19 | 0.20 |
|  | 4 | 0.19 | 0.19 | 0.17 | 0.16 | 0.14 | 0.14 | 0.13 | 0.20 | 0.19 |
| 200 species |  |  |  |  |  |  |  |  |  |  |
| $\mu_{\sigma^2} =$ | -4 | 0.19 | 0.35 | 0.19 | 0.24 | 0.20 | 0.28 | 0.13 | 0.21 | 0.21 |
|  | 0 | 0.17 | 0.09 | 0.17 | 0.21 | 0.23 | 0.19 | 0.11 | 0.21 | 0.21 |
|  | 4 | 0.21 | 0.18 | 0.26 | 0.15 | 0.17 | 0.16 | 0.13 | 0.20 | 0.21 |

**Table 2.** Average posterior breadth for rate variance, trend, and branchwise rate parameters for each simulated trait evolution scenario and tree size

| $\sigma_{\sigma^2}^2 =$ | | rate variance | | | trend | | | branchwise rates | | |
| --- | --- | --- | --- | --- | --- | --- | --- | --- | --- | --- |
|  |  | 0 | 3 | 6 | 0 | 3 | 6 | 0 | 3 | 6 |
| 50 species |  |  |  |  |  |  |  |  |  |  |
| $\mu_{\sigma^2} =$ | -4 | 3.85 | 9.07 | 15.05 | 5.03 | 6.08 | 6.71 | 2.33 | 3.17 | 3.76 |
|  | 0 | 3.65 | 10.07 | 14.82 | 5.92 | 8.26 | 8.28 | 2.29 | 3.41 | 3.90 |
|  | 4 | 4.52 | 8.66 | 14.05 | 10.73 | 10.75 | 10.75 | 3.01 | 3.49 | 3.85 |
| 100 species |  |  |  |  |  |  |  |  |  |  |
| $\mu_{\sigma^2} =$ | -4 | 1.56 | 5.60 | 8.53 | 3.27 | 4.65 | 4.84 | 1.66 | 2.92 | 3.35 |
|  | 0 | 1.91 | 6.45 | 9.01 | 4.31 | 5.27 | 6.01 | 1.87 | 3.10 | 3.42 |
|  | 4 | 1.69 | 6.47 | 8.39 | 7.61 | 8.42 | 7.39 | 2.06 | 3.32 | 3.60 |
| 200 species |  |  |  |  |  |  |  |  |  |  |
| $\mu_{\sigma^2} =$ | -4 | 0.69 | 4.13 | 6.43 | 2.80 | 3.59 | 4.01 | 1.23 | 2.51 | 3.06 |
|  | 0 | 0.62 | 4.23 | 6.21 | 3.39 | 3.99 | 4.06 | 1.18 | 2.72 | 3.23 |
|  | 4 | 0.79 | 3.89 | 6.14 | 4.50 | 5.21 | 5.65 | 1.39 | 2.83 | 3.22 |

**Table 3.**
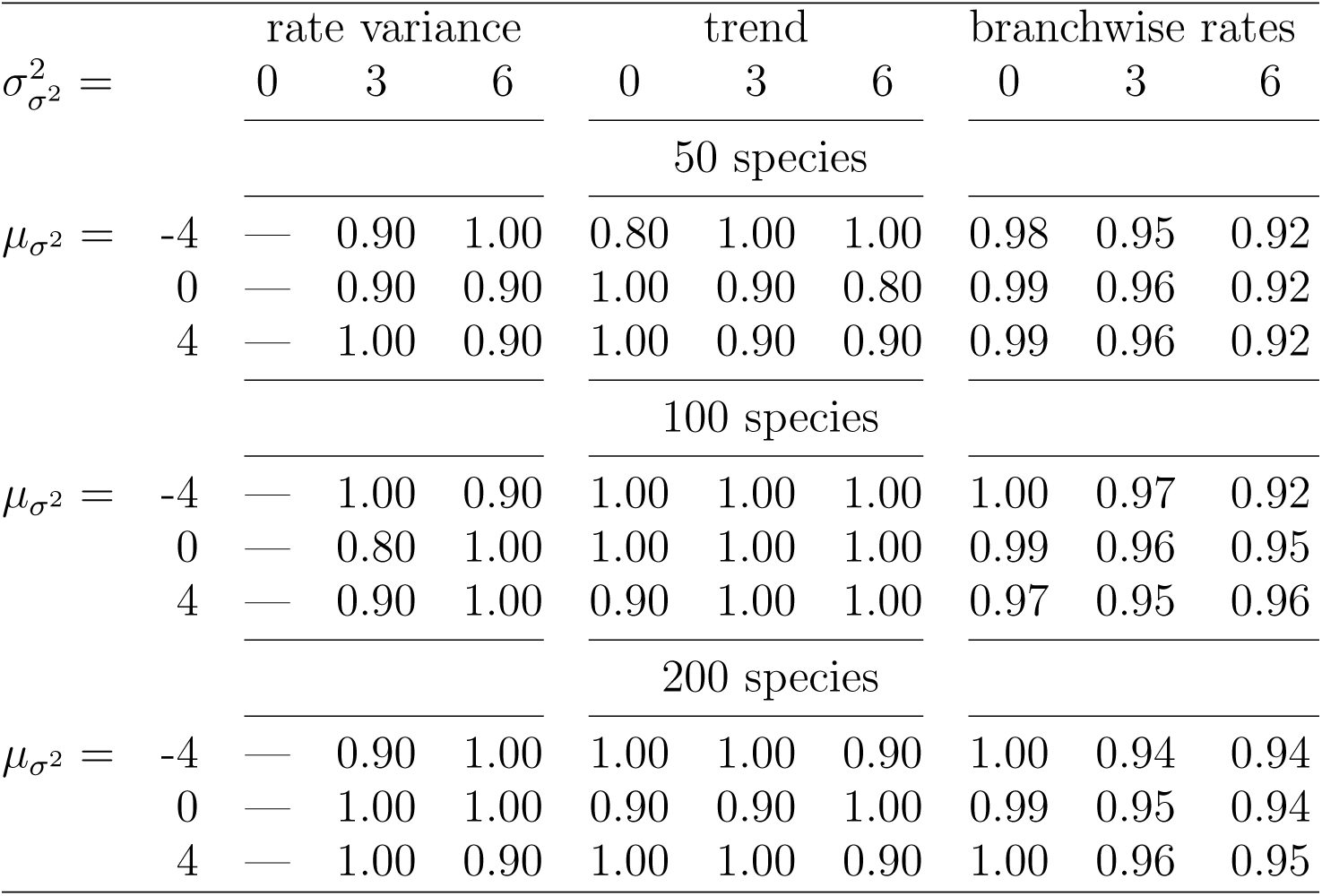
Average posterior coverage for rate variance, trend, and branchwise rate parameters for each simulated trait evolution scenario and tree size

**Figure 1.**
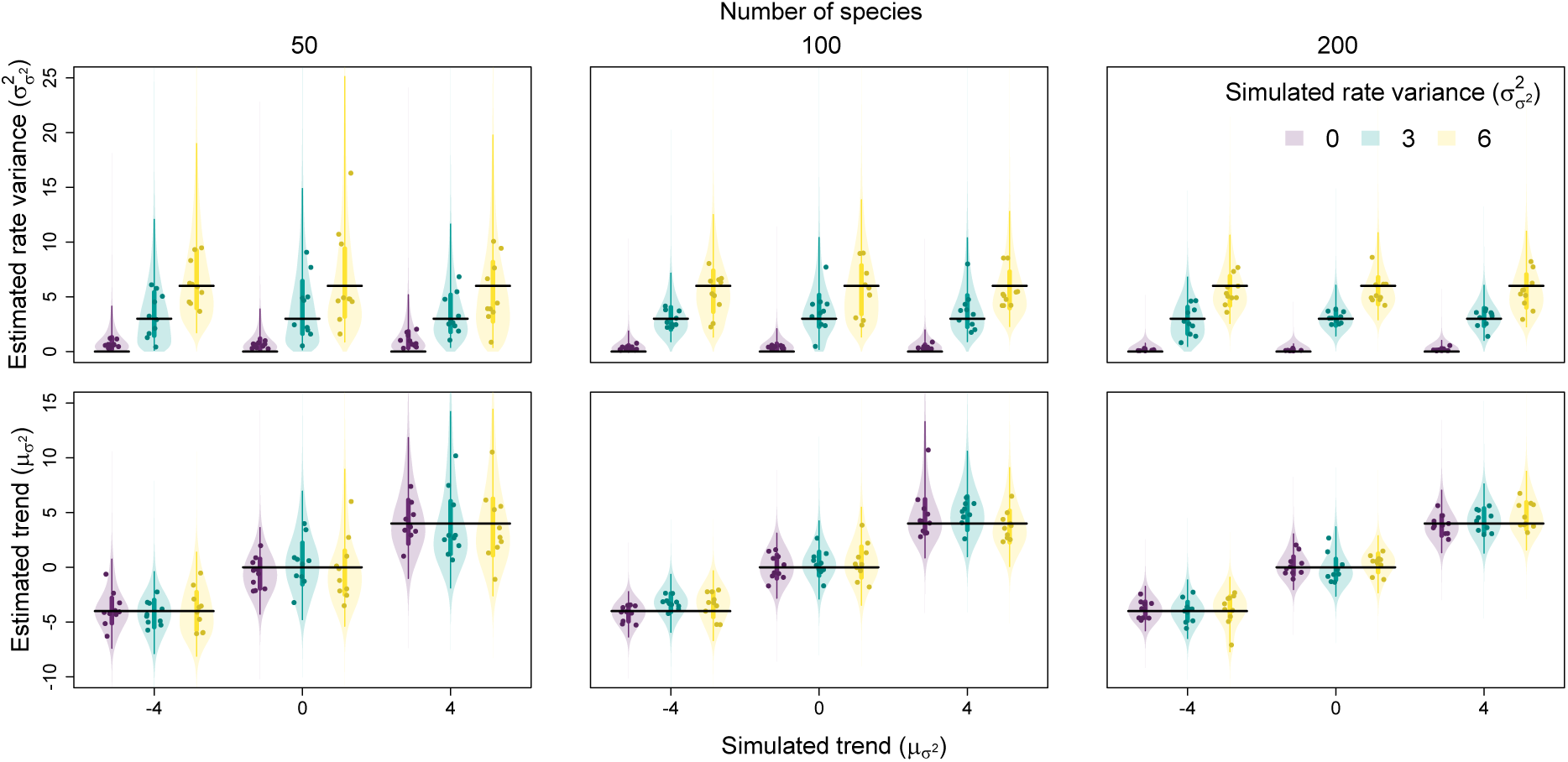
Relationship between simulated and estimated rate variance 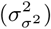 and trend 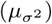 parameters. Each point is the posterior median from a single fit, while the violins are combined posterior distributions from all fits for a given trait evolution scenario. Vertical lines represent the 50% (thicker lines) and 95% credible intervals (thinner lines) of these combined posteriors, while horizontal lines represent positions of true simulated values.

Both the statistical power and error rates of our method were appropriate for detecting trends and significant rate variance. In general, power increased with larger trees, while error rates remained consistent. The ability of SD ratios to identify significant rate variance was particularly impressive, erroneously detecting significant rate variance only once while exhibiting high power (Fig. 2). Decreasing trends were notably easier to detect than increasing trends, particularly on small trees (Fig. 3). Trend error rates consistently remained below *∼*5%, and decreasing trends were never mistaken for increasing trends and vice versa. Higher rate variance seemed to only slightly decrease the power to detect trends. Constraining rate variance to 0 resulted in either worse power or higher error rates for detecting trends, depending on whether trends were decreasing or increasing. As rate variance increased, the power of constrained models to detect decreasing trends dramatically diminished. On the other hand, constrained models detected increasing trends with greater power, at the cost of greatly inflated error rates. Overall, estimating rate variance allows for more sensitive detection of declining trait evolution rates while better safe-guarding against false detection of increasing rates.

**Figure 2.**
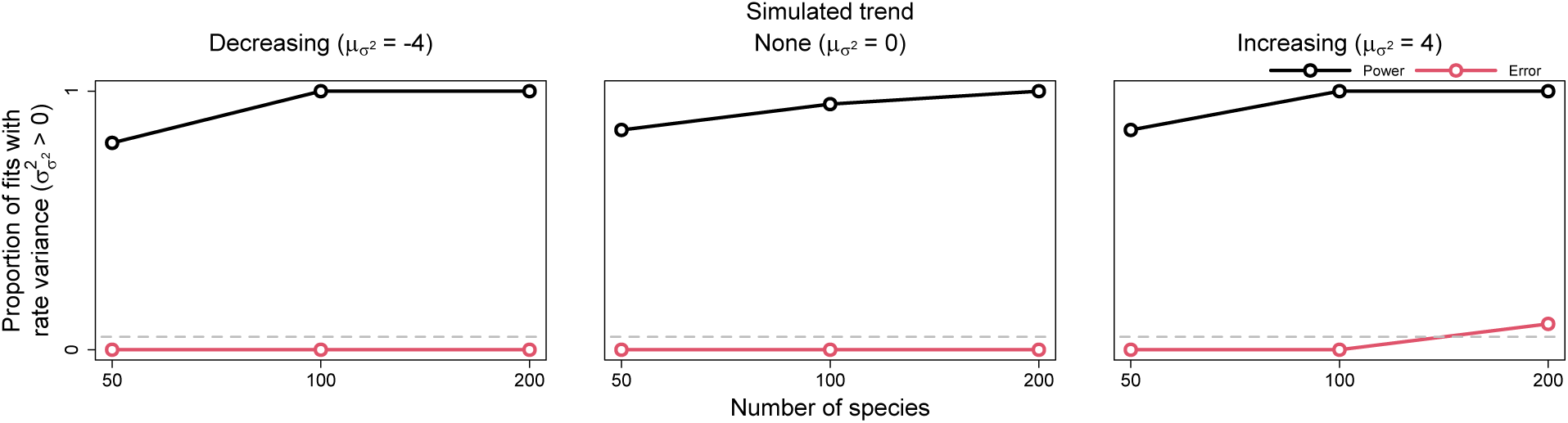
Power and error rates for the rate variance parameter 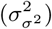. Lines depict changes in the proportion of model fits that correctly showed evidence for rate variance significantly greater than 0 (i.e., power, in black) and incorrectly showed evidence (i.e., error, in red) as a function of tree size.

**Figure 3.**
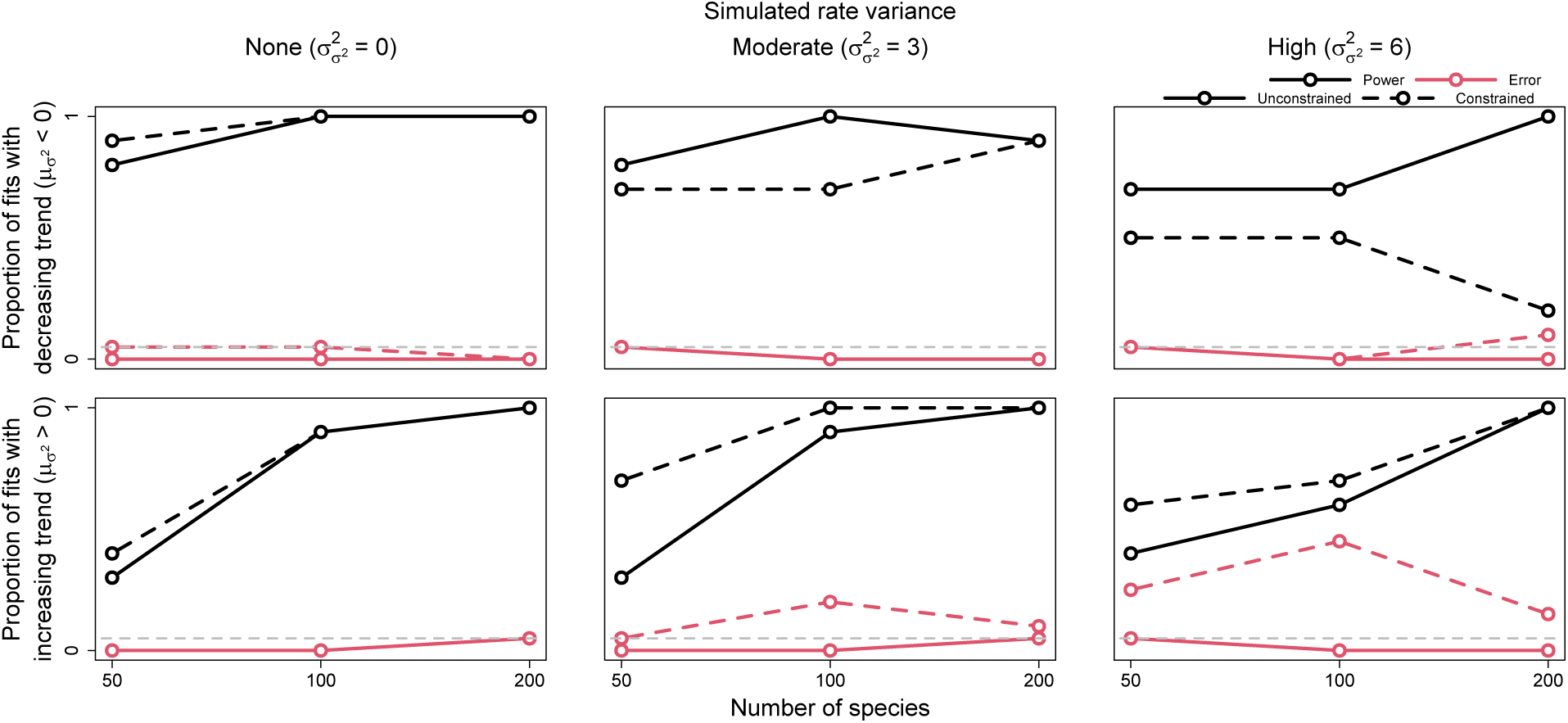
Power and error rates for the trend parameter 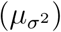. Lines depict changes in the proportion of model fits that correctly showed evidence for trends significantly less and greater than 0 (i.e., power, in black) and incorrectly showed evidence (i.e., error, in red) as a function of tree size. Results are shown for both models allowed to freely estimate rate variance 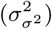 (i.e., unconstrained models, solid lines) and models with rate variance constrained to 0 (i.e., constrained models, dashed lines). The latter models are identical to conventional early/late burst models.

Branchwise rate estimation also generally displayed appropriate coverage, relative accuracy, and statistical testing properties (Tables 1-3, Fig. 4). However, branchwise rate estimates were noticeably biased towards their overall mean (i.e., shrinkage). Linear regressions of median branchwise rate estimates on simulated branchwise rates yield an average slope of about 0.8 (Fig. 5). A similar pattern holds for linear regression of branchwise rate deviations (Fig. S1). Branchwise rate posteriors for simulations with no rate variance exhibited especially high accuracy, precision, and coverage, perhaps due to the increased precision of rate variance posteriors under such trait evolution scenarios. In contrast to other parameters, increasing tree size only slightly decreased posterior breadth for branchwise rates. After accounting for variation in simulated branchwise rate deviations, trait evolution scenario and tree size had little effect on statistical power and error rates for detecting anomalous branchwise rates. Averaging across all fits to simulations with significant rate variance detected, error rates for detecting anomalous rates remained negligible, peaking at around 0.5% for branchwise rate deviations around 0. In fact, this peak only increased to about 5% when we set the significant posterior probability thresholds to 10% and 90% (Fig. S2). The method was somewhat more sensitive to positive than negative deviations, correctly and consistently detecting anomalous rates with deviations more extreme than -4 (1/50th of background rate) or 3 (20 times background rate).

**Figure 4.**
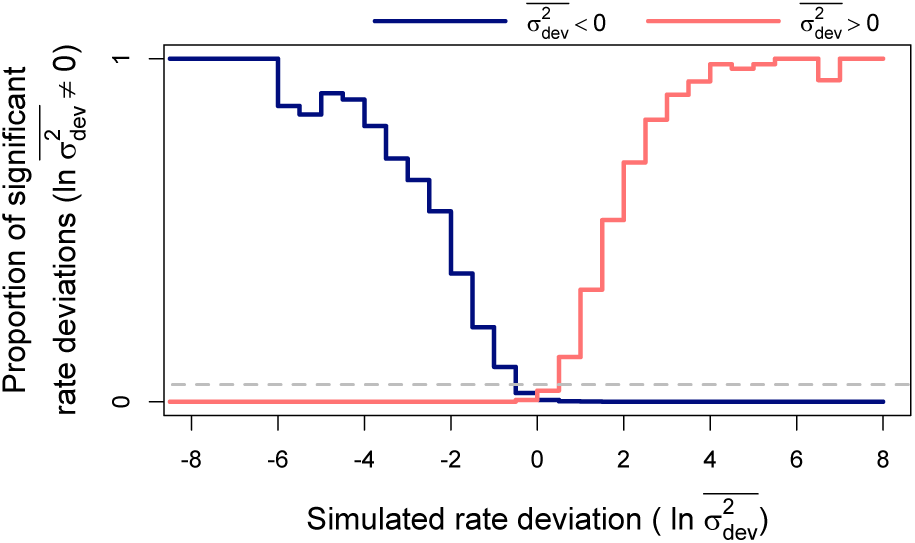
Power and error rates for branchwise rate parameters 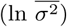. Lines depict changes in proportions of branchwise rates considered anomalously slow (in blue) or fast (in red) as a function of simulated rate deviations 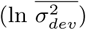. These results combine all fits to simulated data that detected rate variance 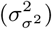 significantly greater than 0. The proportions are equivalent to power when the detected rate deviation is of the same sign as the true, simulated deviation (left of 0 for anomalously slow rates in blue and right for anomalously fast rates in red), and to error rate when the detected and true rate deviations are of opposite signs. Here, significant rate deviations for simulated rate deviations that are exactly 0 are considered errors regardless of sign.

**Figure 5.**
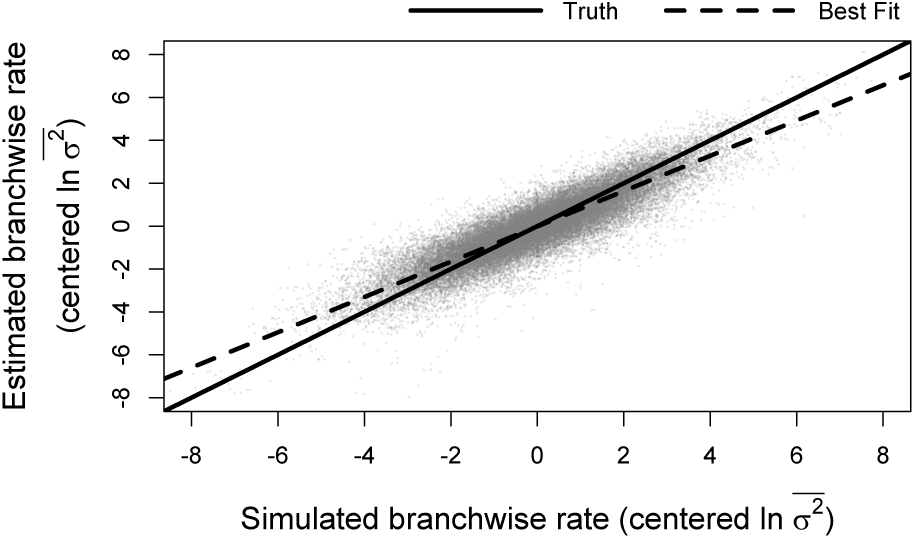
Relationship between simulated and estimated branchwise rate parameters 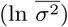. For each simulation and posterior sample, branchwise rates were first centered by subtracting their mean. We estimated centered branchwise rates by taking the median of the centered posterior samples. The solid line represents the position of the true centered branchwise rates, while the shallower, dashed line represents the observed line of best fit for these data.

### Empirical Example

Overall, our model suggests that rates of body size evolution among extant cetaceans have generally slowed down over time, with considerable divergence in rates of body size evolution among key clades (Fig. 6). We found marginally significant support for a decreasing trend in rates over time, with rates declining by about 7% every million years (95% credible interval (CI): 0 - 15% decrease, posterior probability (PP) of increasing trend: 2.5%). We also infer a moderate rate variance of about 0.06 per million years (CI: 0.01 - 0.22, SD ratio: 0.14). Combining these two results, changes in body size evolution rates over a million year time interval are expected to range from a 50% decrease to 60% *increase* for any particular lineage (Fig. 7).

**Figure 6.**
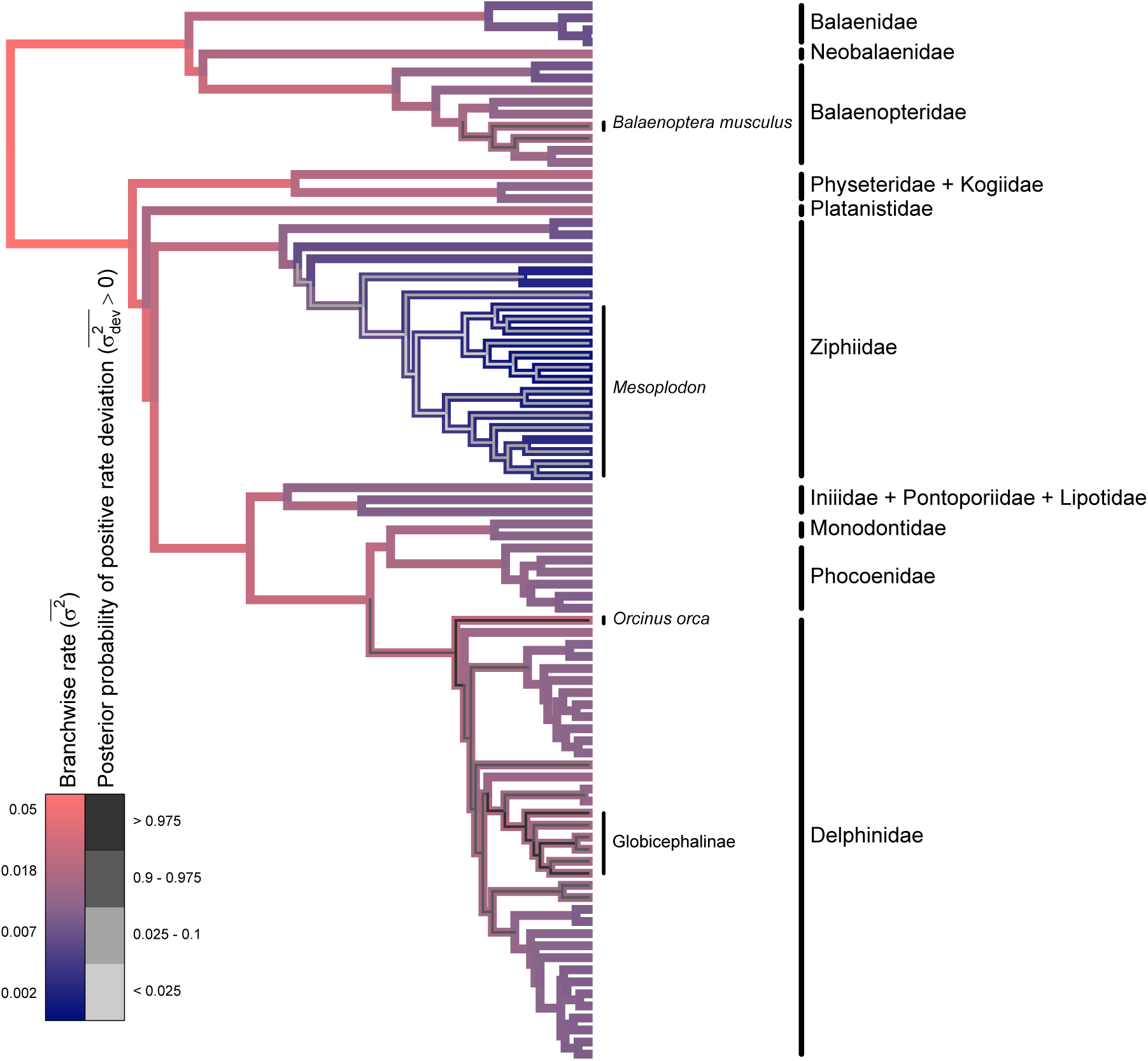
Phylogram of model results for cetacean body size data. Branch colors represent median posterior estimates of branchwise rates 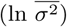 of body size evolution, with slower and faster rates in blue and red, respectively. The thinner, inset colors represent the posterior probability that a branchwise rate is anomalously fast according to its rate deviation 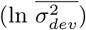, with lower and higher posterior probabilities in light and dark gray, respectively.

**Figure 7.**
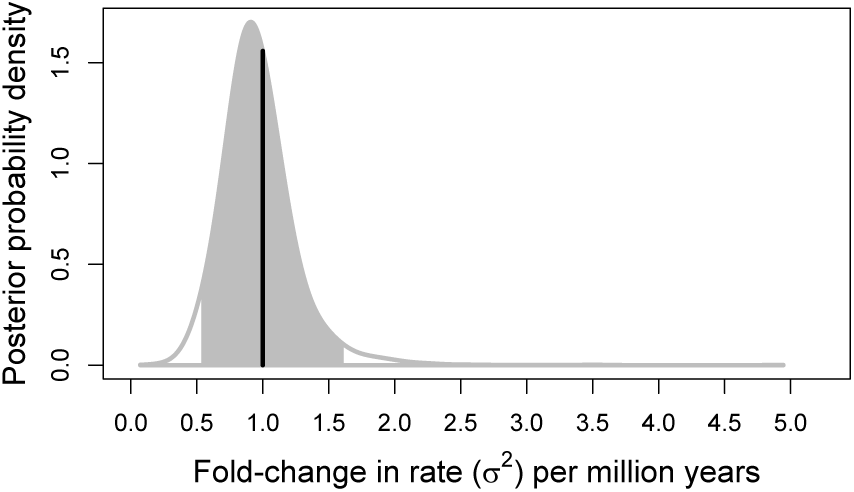
The posterior probability distribution of fold-changes in cetacean body size evolution rates (*σ*^2^) per 1 million years. This distribution is given by 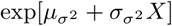, where *X* is a random variable drawn from a standard normal distribution. The gray filled-in portion represent the 95% credible interval, while the vertical line represents the starting rate of 1.

We also identify a few regions of the cetacean phylogeny where rates of body size evolution seem to be especially low or high. After detrending rates, rates of body size evolution in the beaked whale genus *Mesoplodon* are about 34% slower than the background rate (CI: 13 - 77%, PP of positive rate deviation: *<* 1%). On the other end of the spectrum, some oceanic dolphin lineages exhibit unusually rapid body size evolution rates. In particular, pilot whales and allies (subfamily globicephalinae) and the orca (*Orcinus orca*) lineage exhibit body size evolution rates about 160% (CI: 10 - 900%, PP: 99%) and 200% (CI: 20 - 1300, PP: 99%) higher than the background rate, respectively. In fact, oceanic dolphins as a whole exhibit a marginally significant increase in body size evolution rates, even after excluding the pilot whale subfamily and orca lineage (CI: 90 - 300% background rate, PP: 95%). The blue whale (*Balanoptera musculus*) lineage also exhibits a marginally significant increase in body size evolution rate, about 140% (95% CI: -10 - 1000%, PP: 95%) higher than the background rate.

Under the model with rate variance constrained to 0, rates of body size evolution decrease by only about 4% every million years (95% CI: -1 - 10%, PP of increasing trend: 7.3%). While only a slight difference, the trend parameter estimated under the full model yields a marginally significant, two-tailed “p-value” of *∼*5%, while the constrained model yields a decidedly insignificant “p-value” of *∼*15%. This is reflected in a conventional sample-size corrected Akaike Information Criterion (AICc) comparison between simple BM and EB models of trait evolution fitted via maximum likelihood using the R package *geiger* (version 2.0.7; Pennell et al., 2014). In this case, a simple BM model receives nearly twice the AICc weight of an EB model (65% vs. 35%).

## Discussion

Here we implemented a novel data-driven method, *evorates*, for modeling stochastic, incremental variation in trait evolution rates. Part of the power of *evorates* is its ability to infer trait evolution rate variation independent of an *a priori* hypothesis on what factors influence rates. This allows for detailed, hypothesis-free exploration of trait evolution rate variation across time and taxa. Researchers may use such results to generate and refine hypotheses regarding what factors have influenced trait evolution rates across the tree of life (e.g., Uyeda et al., 2018). Overall, *evorates* performs well on simulated data, recovering accurate parameter estimates and exhibiting appropriate statistical power and error rates when testing the significance of parameters. Further, the method shows great promise for empirical macroevolutionary research, offering novel insights into the dynamics of cetacean body size evolution–a notably well-studied system (e.g., Slater et al., 2010, Pyenson and Sponberg, 2011 Montgomery et al., 2013, Slater and Pennell, 2014, Slater et al., 2017, Sander et al., 2021). The results of our study also builds on previous work in demonstrating that estimating time-independent rate heterogeneity is critical for accurately quantifying temporal dynamics in trait evolution rates (Slater and Pennell, 2014). This finding has consequences for how EBs/LBs of trait evolution are practically identified and conceptually defined.

The simulation study results showcases *evorate’s* ability to recover accurate parameter estimates across a range of tree sizes. Despite the high uncertainty of rate variance estimates under some trait evolution scenarios, rate heterogeneity could still be correctly detected about 90% of the time with an error rate substantially lower than 5%. Compared to conventional EB/LB models, our method can detect decreasing trends in trait evolution rates with greater sensitivity and detect increasing trends with greater robustness. Notably, traits evolving with exponentially increasing rates on an ultrametric phylogeny (i.e., an LB model) exhibit the same probability distribution expected under a single-peak Ornstein-Uhlenbeck (OU) model, where traits evolve towards some optimum at a constant rate (Blomberg et al., 2003). The fact that autocorrelated rate heterogeneity increases support for such models has not been noted before to our knowledge, and offers an alternative explanation for frequently observed support for OU models in comparative data (e.g., Harmon et al., 2010; see also Cooper et al., 2016; Landis and Schraiber, 2017). Despite their mathematical similarities, LB, OU, and our new models have distinct biological interpretations regarding the importance of rate heterogeneity and selective forces in shaping the patterns of trait diversity within clades.

Interestingly, closer inspection of our simulation study results suggest that, in the presence of rate heterogeneity, models with rate variance constrained to 0 (i.e., conventional EB/LB models) estimate trend parameters corresponding to changes in average trait evolution rates over time. On the other hand, unconstrained *evorates* models estimate trend parameters corresponding to changes in median trait evolution rates over time, essentially determining whether most lineages in a clade exhibit rate decreases or increases (Figs. S19-21; Tables S5-7). Counterintuitively, when the trend parameter is only weakly negative relative to rate variance 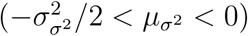, it is possible for a majority of lineages within a clade to exhibit declining trait evolution rates (i.e., an EB according to *evorates*) while rates averaged across the entire clade increase over time (i.e., an LB according to conventional methods). This occurs because rates evolve in a right-skewed manner under our model–in other words, a few anomalous lineages/subclades tend to evolve extremely high trait evolution rates in spite of declining rates among most other lineages, driving up a clade’s overall average rate (Fig. S17-18). We note that *evorates* still returns estimates of average changes in trait evolution rates per unit time via a simple parameter transformation 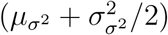. We choose to focus on the majority-based definition of EBs/LBs since, by accounting for anomalous lineages/subclades exhibiting unusual rates, this definition better matches many macroevolutionary biologists’ intuitive definition of EBs (Lloyd et al., 2012; Slater and Pennell, 2014; Benson et al., 2014; Hopkins and Smith, 2015; Wright, 2017; Puttick, 2018).

Our empirical example with cetacean body size directly demonstrates the practical importance of these nuances in defining EB/LB dynamics. We find substantial evidence that body size evolution has slowed down in most cetacean lineages, despite the presence of “outlier” lineages exhibiting relatively rapid rates. Indeed, we find little evidence for a decline in body size evolution rates averaged across the clade (CI: 12% decrease - 5% increase in average rate per million years, posterior probability of increasing trend: 16%). This broadly agrees with previous research, but *evorates* is able to offer novel insights and contextualize prior results by explicitly estimating branchwise rates in addition to overall trends (Slater and Pennell, 2014; Sander et al., 2021). For example, Slater and Pennell (2014) identified the orca and pilot whale lineages as outlier lineages exhibiting especially rapid rates of body size evolution. Our method recapitulates these findings while suggesting oceanic dolphins as a whole represent a relatively recent burst of body size evolution that has largely masked signals of an earlier burst towards the base of the clade. Such findings more generally agree with recent suggestions that bursts of trait evolution may be common but not limited to the base of “major” clades. This is likely due, in part, to major clades being arbitrarily designated based on taxonomic rank (Puttick, 2018). Alternatively, some propose that EBs may be hierarchical, with major clades exhibiting repeated bouts of rapid trait diversification as competing, closely-related lineages partition niche space more finely over time (Slater and Friscia, 2019). Ultimately, we are optimistic that *evorates* may be better able to resolve how frequently bursts of trait evolution–early or not–occur across the tree of life compared to more conventional methods.

The shrinkage of branchwise rates, whereby rate estimates are biased towards their overall mean, is presumably due to the assumption that rates are autocorrelated under our model. Because of this, rate estimates are partially informed by the rates in closely-related lineages, particularly when closely-related lineages are better sampled (i.e., more related to taxa with sampled trait values and/or consisting of many short branch lengths). This “diffusion” of rates across the phylogeny appears to cause under- and overestimation of unusually high and low rates, respectively. Fortunately, this renders *evorates* conservative in terms of identifying anomalous trait evolution rates, safeguarding against erroneous conclusions. In general, we view this behavior as a good compromise between model flexibility and robustness, allowing *evorates* to infer rate variation while avoiding ascribing significance to noise in data. We note that rate variance estimates under our model are largely unbiased, such that branchwise rates in a typical posterior sample should be as variable as the true rates. Thus, taking the joint distribution of branchwise rates into account by analyzing distributions of *differences* between rates, rather than just assessing marginal distributions of rates, appears important in accurately interpreting results under our model. In any case, despite this shrinkage phenomenon, the statistical power to identify overall rate heterogeneity and anomalous rates with *evorates* appears comparable to that of previous data-driven methods (Eastman et al., 2011).

*Evorates* is one of several recently developed methods that also estimate unique trait evolution rates for each branch in a phylogeny but assume an alternative mode of rate change (May and Moore, 2020; Fisher et al., 2021). These other methods assume that branchwise rates are independently distributed according to a log-normal distribution. The method we develop here differs from these “independent rate” (IR) models in assuming that rates evolve gradually and are thus phylogenetically autocorrelated (see also Revell, 2021). Theoretically, trait evolution rates should exhibit some degree of phylogenetic autocorrelation given that many factors hypothesized to affect trait evolution rates themselves exhibit phylogenetic autocorrelation. Indeed, a recent study found evidence for autocorrelation of trait evolution rates in a few vertebrate clades (Sakamoto and Venditti, 2018), and autocorrelation has also been found in lineage diversification (Savolaine et al., 2002; Caron and Pie, 2020) and molecular substitution rates (Lepage et al., 2006; Tao et al., 2019). Notably, there is also no known rate evolution process that would produce independent, log-normally distributed branchwise rates (Lepage et al., 2006, 2007). However, IR models could outperform “autocorrelated rate” (AR) models in some instances due to their tremendous flexibility in modeling how rates vary over time and phylogenies. In general, we expect that IR models will perform best in cases with many traits and/or non-ultrametric trees, where the flexibility of the model can be tempered by rich information content in the data. More work testing for rate autocorrelation or lack thereof in continuous trait data is needed as methods for inferring trait evolution rate variation become more complex.

Revell (2021) independently developed a method, *multirateBM*, based on a model similar to the one we introduce here, though *evorates* offers several key advantages. In particular, the maximum likelihood (ML) implementation of *multirateBM* renders it impossible to estimate rate variance. To do so, one would need to analytically marginalize over uncertainty in branchwise rates. Here, we circumvent this issue by using Bayesian inference to numerically integrate over uncertainty in branchwise rates. This is analogous to how ML implementations of mixed effect models analytically marginalize over uncertainty in random effects, while Bayesian implementations of the same models sample random effects (Browne and Draper, 2006). Indeed, ML implementations of mixed effect models that treat random effects as parameters would be unable to estimate random effect variances due to the very same reasons *multirateBM* cannot estimate rate variance. Additionally, our model has the added advantage of accommodating both trends in rates and uncertainty in tip trait values. Lastly, we implement procedures to test the significance of rate heterogeneity, trends, and anomalous trait evolution rates. While *multirateBM* offers a quick and convenient means for comparative data exploration, our new method allows for more rigorous quantification and analysis of rate evolutionary processes and patterns from comparative data.

There are a number of ways the *evorates* might be improved or expanded. Assuming that trait evolution rates for different traits are correlated with one another, using data on multiple traits could improve inference of both the rate evolution process and branchwise rate parameters (May and Moore, 2020). Another promising future direction is integration of *evorates* with hypothesis-driven methods. This could be done *post hoc* by applying phylogenetic linear regression to “tip rates” estimated under the model (e.g., Rabosky and Huang, 2016) or analyzing distributions of branchwise rates associated with ancestral states estimated via stochastic character maps (Revell, 2013; but see May and Moore, 2020). Alternatively, one could explicitly model rates as the product of both a stochastic rate evolution process and a deterministic function of some factor of interest. We have already taken steps towards this model extension in our current implementation by allowing rates to change as a deterministic function of time. Lastly, despite our focus on gradually changing rates, trait evolution rates might also exhibit sudden shifts of large magnitude (“jumps”) or short-lived fluctuations (“pulses”) in response to factors with particularly strong influence on rates. It would be ideal–but difficult–to model rates as evolving gradually, while potentially undergoing sudden jumps or pulses (e.g., Lartillot et al., 2016). An alternative strategy is developing methods to compare the fit of a model like ours against more conventional data-driven models whereby rates jump or even L’evy models whereby rates pulse (Landis et al., 2013). Assessing when and whether comparative data can distinguish between different modes of rate change will be important for future research on the dynamics of trait evolution.

### Conclusion

Here, we introduced *evorates*, a method that models gradual change, rather than abrupt shifts, in continuous trait evolution rates from comparative data. Unlike nearly all other comparative methods for inferring rate variation, *evorates* goes beyond identifying lineages exhibiting anomalous rates by also estimating the process by which rates themselves evolve. Although there are many potential modes of rate variation over time and phylogenies, our model estimates rate evolution processes as the product of two parameters: one controlling how quickly rates accumulate random variation, and another determining whether rates tend to decrease or increase over time. The resulting method returns accurate estimates of evolutionary processes and provides a flexible and intuitive means of detecting and analyzing trait evolution rate variation. Looking forward, *evorates* has tremendous potential for improvement and elaboration, and we are optimistic that the future of macroevolutionary biology will benefit from increased focus not only on how traits evolve, but how the rates of trait evolution themselves evolve over time and taxa.

## Acknowledgements

We would like to thank Robert Lanfear and Bob Week for their feedback on mathematical reasoning and intuition underlying our method. We are also grateful to Graham Slater for offering valuable insights and advice regarding our empirical example. Lastly, we would like to thank members of the Bradburd and Weber labs at Michigan State University for general feedback on project development and writing.

## Supplementary Material

Data available from the Dryad Digital Repository: http://dx.doi.org/10.5061/dryad.[NNNN] (coming soon, but all files are available upon request to B.S.M. at; current version of online appendix is available as supplementary material on bioRxiv). The current version of the *evorates* R package is available at the GitHub repository: https://github.com/bstaggmartin/evorates.

