## Supplementary material for "Modeling the Evolution of Rates of Continuous Trait Evolution": Online Appendix

### Supplemental Tables and Figures

Table S1. *Cetacean body length data and associated references used for empirical example*

| species | length (m) | reference |
|---|---|---|
| *Balaena mysticetus* | 18.0 | Slater et al., 2010 |
| *Balaenoptera acutorostrata* | 10.7 | Slater et al., 2010 |
| *Balaenoptera bonaerensis* | 10.2 | Konishi et al., 2008 |
| *Balaenoptera borealis* | 16.1 | Slater et al., 2010 |
| *Balaenoptera edeni* | 15.4 | Slater et al., 2010 |
| *Balaenoptera musculus* | 33.6 | Slater et al., 2010 |
| *Balaenoptera omurai* | 10.7 | Slater et al., 2010 |
| *Balaenoptera physalus* | 21.2 | Slater et al., 2010 |
| *Berardius arnuxii* | 8.9 | Slater et al., 2010 |
| *Berardius bairdii* | 12.0 | Slater et al., 2010 |
| *Caperea marginata* | 6.2 | Slater et al., 2010 |
| *Cephalorhynchus commersoni* | 1.5 | Slater et al., 2010 |
| *Cephalorhynchus eutropia* | 1.5 | Molina and Oporto, 1993 |
| *Cephalorhynchus heavisidii* | 1.7 | Slater et al., 2010 |
| *Cephalorhynchus hectori* | 1.5 | Slater et al., 2010 |
| *Delphinapterus leucas* | 3.8 | Slater et al., 2010 |
| *Delphinus capensis* | 2.5 | Plön et al., 2012 |
| *Delphinus delphis* | 2.3 | Slater et al., 2010 |
| *Eschrichtius robustus* | 14.6 | Slater et al., 2010 |
| *Eubalaena australis* | 13.9 | Slater et al., 2010 |
| *Eubalaena glacialis* | 13.7 | Slater et al., 2010 |
| *Eubalaena japonica* | 17.4 | Fortune et al., 2021 |
| *Feresa attenuata* | 2.4 | Slater et al., 2010 |
| *Globicephala macrorhynchus* | 4.8 | Slater et al., 2010 |
| *Globicephala melas* | 5.1 | Slater et al., 2010 |
| *Grampus griseus* | 3.7 | Slater et al., 2010 |
| *Hyperoodon ampullatus* | 7.9 | Slater et al., 2010 |
| *Hyperoodon planifrons* | 7.5 | Slater et al., 2010 |
| *Indopacetus pacificus* | 7.2 | Slater et al., 2010 |
| *Inia geoffrensis* | 2.0 | Slater et al., 2010 |
| *Kogia breviceps* | 3.4 | Slater et al., 2010 |
| *Kogia sima* | 2.4 | Slater et al., 2010 |
| *Lagenodelphis hosei* | 2.6 | Slater et al., 2010 |
| *Lagenorhynchus albirostris* | 3.0 | Slater et al., 2010 |
| *Leucopleurus acutus* | 2.4 | Slater et al., 2010 |
| *Lipotes vexillifer* | 2.0 | Slater et al., 2010 |
| *Lissodelphis borealis* | 2.3 | Slater et al., 2010 |
| *Lissodelphis peronii* | 2.3 | Baker, 1981 |

| species | length (m) | reference |
|---|---|---|
| *Megaptera novaeangliae* | 18.0 | Slater et al., 2010 |
| *Mesoplodon bidens* | 5.1 | Slater et al., 2010 |
| *Mesoplodon bowdoini* | 4.5 | Slater et al., 2010 |
| *Mesoplodon carlhubbsi* | 5.3 | Mead et al., 1982 |
| *Mesoplodon densirostris* | 4.7 | Slater et al., 2010 |
| *Mesoplodon europaeus* | 5.2 | Slater et al., 2010 |
| *Mesoplodon ginkgodens* | 4.9 | Slater et al., 2010 |
| *Mesoplodon grayi* | 5.3 | Slater et al., 2010 |
| *Mesoplodon hectori* | 4.4 | Slater et al., 2010 |
| *Mesoplodon hotaula* | 4.8 | Dalebout et al., 2014 |
| *Mesoplodon layardii* | 6.2 | Slater et al., 2010 |
| *Mesoplodon mirus* | 5.1 | Slater et al., 2010 |
| *Mesoplodon perrini* | 4.4 | Dalebout et al., 2002 |
| *Mesoplodon peruvianus* | 3.7[a] | Reyes et al., 1991 |
| *Mesoplodon stejnegeri* | 5.7 | Slater et al., 2010 |
| *Mesoplodon traversii* | 5.3 | Thompson et al., 2012 |
| *Monodon monoceros* | 4.3 | Slater et al., 2010 |
| *Neophocaena phocaenoides* | 1.4 | Slater et al., 2010 |
| *Orcaella brevirostris* | 2.2 | Slater et al., 2010 |
| *Orcaella heinsohni* | 2.2 | Arnold and Heinsohn, 1996 |
| *Orcinus orca* | 7.9 | Slater et al., 2010 |
| *Peponocephala electra* | 2.8 | Lodi et al., 1990 |
| *Phocoena dioptrica* | 1.9 | Slater et al., 2010 |
| *Phocoena phocoena* | 1.9 | Slater et al., 2010 |
| *Phocoena sinus* | 1.1 | Slater et al., 2010 |
| *Phocoena spinipinnis* | 1.7 | Slater et al., 2010 |
| *Phocoenoides dalli* | 1.9 | Slater et al., 2010 |
| *Physeter macrocephalus* | 11.0 | Slater et al., 2010 |
| *Platanista gangetica* | 2.5 | Slater et al., 2010 |
| *Pontoporia blainvillii* | 1.5 | Slater et al., 2010 |
| *Pseudorca crassidens* | 5.1 | Slater et al., 2010 |
| *Sagmatias australis* | 2.1 | Slater et al., 2010 |
| *Sagmatias cruciger* | 1.8 | Slater et al., 2010 |
| *Sagmatias obliquidens* | 2.4 | Slater et al., 2010 |
| *Sagmatias obscurus* | 1.9 | Slater et al., 2010 |
| *Sotalia fluviatilis* | 1.5 | Slater et al., 2010 |
| *Sotalia guianensis* | 2.1 | Barros, 1991 |
| *Sousa chinensis* | 2.4 | Slater et al., 2010 |
| *Sousa teuszii* | 2.5 | Jefferson and Rosenbaum, 2014 |
| *Stenella attenuata* | 2.1 | Slater et al., 2010 |
| *Stenella clymene* | 1.9 | Slater et al., 2010 |
| *Stenella coeruleoalba* | 2.3 | Slater et al., 2010 |
| *Stenella frontalis* | 2.1 | Slater et al., 2010 |

| species | length (m) | reference |
|---|---|---|
| *Stenella longirostris* | 2.0 | Slater et al., 2010 |
| *Steno bredanensis* | 2.6 | Slater et al., 2010 |
| *Tasmacetus shepherdi* | 6.5 | Slater et al., 2010 |
| *Tursiops aduncus* | 2.1 | Slater et al., 2010 |
| *Tursiops australis* | $2.8^b$ | Charlton-Robb et al., 2011 |
| *Tursiops truncatus* | 2.4 | Slater et al., 2010 |
| *Ziphius cavirostris* | 6.4 | Slater et al., 2010 |

[a]from male specimen since no mature females were measured

[b]sex not reported

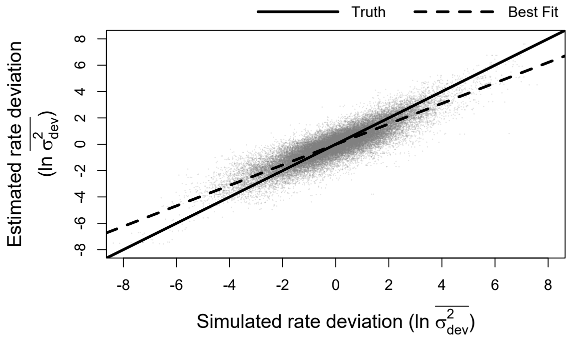


Figure S1. Relationship between simulated and estimated branchwise rate deviation parameters (ln $\overline{\sigma^2_{dev}}$). The solid line represents the position of the true branchwise rate deviations, while the shallower, dashed line represents the observed line of best fit for these data.

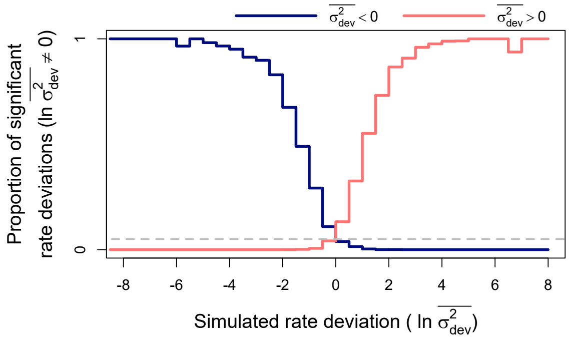


Figure S2. Power and error rates for branchwise rate parameters (ln $\overline{\sigma^2}$) under relaxed significance thresholds (posterior probability $< 0.1$ or $> 0.9$). Lines depict changes in proportions of branchwise rates considered anomalously slow (in blue) or fast (in red) as a function of simulated rate deviations (ln $\overline{\sigma^2_{dev}}$). These results combine all fits to simulated data that detected rate variance ($\sigma^2_{\sigma^2}$) significantly greater than 0. The proportions are equivalent to power when the detected rate deviation is of the same sign as the true, simulated deviation (left of 0 for anomalously slow rates in blue and right for anomalously fast rates in red), and to error rate when the detected and true rate deviations are of opposite signs. Here, significant rate deviations for simulated rate deviations that are exactly 0 are considered errors regardless of sign.

### Approximating Geometric Brownian Motion Time-Averages

Our model seeks to model rates ($\sigma^2$) as "evolving" under a trended Geometric Brownian Motion (GBM)-like process, whereby the natural log of rates evolve in a trended Brownian Motion (BM)-like manner. Unfortunately, this requires an expression for the probability distribution of GBM time-averages along each branch in the phylogeny. Expressions for such distributions are infamously intractable, necessitating approximate solutions (Dufresne, 2004; Lepage et al., 2007). For our model, we use a multivariate log-normal approximation to model rate time-averages along each branch (branchwise averages, $\bar{\sigma^2}$) based on two observations. First, as the rate variance parameter ($\sigma^2_{\sigma^2}$) approaches 0, rates ($\sigma^2$) will converge to following a simple exponential function with respect to time, $\sigma^2 = \sigma_0^2 \exp[\mu_{\sigma^2} t]$, where $\sigma_0^2$ is the starting rate, $\mu_{\sigma^2}$ is the trend, and $t$ is time. In this case, the branchwise averages can be derived through integration and are equivalent to the time-averaged rates expected under a conventional "early/late burst" (EB/LB) model (Blomberg et al., 2003). Second, over short amounts of time and/or with low rate variance, the arithmetic and geometric time-averages of a GBM process approach one another. The geometric time-average of a GBM process is simply the exponentiated arithmetic time-average of the GBM process on the natural log scale, which has a straight-forward and tractable log-normal distribution (Devreese et al., 2010). Thus, assuming that branch lengths in a phylogeny are typically short and rate variance is relatively low, we can approximate the distribution of the natural log of branchwise averages by adding multivariate normal "noise", $\gamma$, to the natural log of branchwise averages expected under a conventional EB/LB model, $\beta$. In other words:

$$\ln(\overline{\sigma^2}) \approx \beta + \gamma \tag{1}$$

$$\beta = \ln(\sigma_0^2) + \begin{cases} 0 & \text{if } \mu_{\sigma^2} = 0 \\ \ln(|\exp[\mu_{\sigma^2}\tau_2] - \exp[\mu_{\sigma^2}\tau_1]|) - \ln(|\mu_{\sigma^2}|) - \ln(t) & \text{if } \mu_{\sigma^2} \neq 0 \end{cases} \tag{2}$$

$$\gamma \sim MVN(0, \sigma^2_{\sigma^2} D) \tag{3}$$

as in the main text. Here, $t$ is a vector of branch lengths, $\tau_1$ and $\tau_2$ are vectors of the start and end times of each branch (i.e., $\tau_2 - \tau_1 = t$), and $D$ is the variance-covariance matrix of branchwise averages for a value evolving under an untrended BM process on a phylogeny. Let $\bar{x}$ and $t$ be vectors of time-averaged trait values and edge lengths, respectively, for three edges: two sister edges, $i$ and $j$, with ancestral edge, $k$. If traits evolve under an untrended BM process and the ancestral trait value of $k$ is fixed, the variances of $\bar{x_i}$ and $\bar{x_j}$ are $t_i/3 + t_k$ and $t_j/3 + t_k$, respectively. The covariance between $\bar{x_i}$ and $\bar{x_j}$ is simply $t_k$, and the covariances between either $\bar{x_i}$ or $\bar{x_j}$ and $\bar{x_k}$ is $\bar{x_k}/2$ (Devreese et al., 2010). From this, we can derive an expression for the variance-covariance matrix of branchwise averages given an arbitrary phylogeny, as shown in the main text:

$$D_{i,j} = \sum_{k \in \mathrm{anc}(i,j)} t_k - \begin{cases} 2t_i/3 & \text{if } i = j \\ t_i/2 & \text{if } i \in \mathrm{anc}(j,j) \\ t_j/2 & \text{if } j \in \mathrm{anc}(i,i) \\ 0 & \text{if } i \neq j, i \notin \mathrm{anc}(j,j), j \notin \mathrm{anc}(i,i) \end{cases} \tag{4}$$

While this multivariate log-normal approximation is rough, we demonstrate here that it is largely sufficient for our purposes. Notably, we are not the first to approximate GBM time-averages using log-normal distributions in the context of comparative phylogenetics (Welch and Waxman, 2008). There are two other tractable strategies for approximating these distributions given in the comparative phylogenetics literature. Both of these strategies use the fact that values at the nodes of a phylogeny evolving under a GBM process follow an exact multivariate log-normal distribution, and instead focus on estimating nodewise values. Branchwise averages are then approximated by either averaging ancestral and descendant nodes for each edge (e.g., Thorne et al., 1998) or via the maximum likelihood estimate of branchwise averages given the ancestral and descendant nodes (e.g., Lartillot and Poujol, 2011; Revell, 2021). We term these strategies "endpoint averaging" and "endpoint integration", respectively. We prefer the log-normal approximation due to its convenient formulation and direct focus on estimating branchwise, rather than nodewise, quantities. In the spirit of thoroughness, however, we conducted three simulation experiments to investigate the relative performance of these different approximation strategies.

We first conducted a simple experiment where we simulated 100,000 GBM time-averages on the natural log scale under each approximation strategy. We also estimated a "true" branchwise average distribution for comparison by simulating 100,000 fine-grained GBM sample paths (1,000 time points) and taking the natural log of each sample path's average. We repeated these simulations for each combination of trend ($\mu_{\sigma^2}$) and rate variance ($\sigma^2_{\sigma^2}$) parameter values used in the main text's simulation study (Fig. S3). All simulations were standardized to occur over a time interval of 1, just as each phylogeny in our simulation study was rescaled to have a total height of 1. The results below thus represent how "off" each approximation would be for a single branch spanning the entire height of a phylogeny in our simuation study. The log-normal approximation notably lacks a right skew characteristic of the true distribution and other approximations. The log-normal approximation also appears to overestimate the variance of branchwise averages when trends are decreasing and underestimates variance when trends are increasing, particularly with high rate variance. On the other hand, the endpoint average approximation exhibits notable upward bias and consistently underestimates branchwise average variance. Additionally, this approximation fails to converge to the correct branchwise average when rate variance is 0. Lastly, the endpoint integration approximation exhibits no notable bias but underestimates branchwise average variance in the case of no or decreasing trends. The accuracy of branchwise average variance under the log-normal approximation might be improved by adapting the Fenton-Wilkinson approximation of log-normal sums for GBM processes (Safak and Safak, 2002), but we did not explore this here.

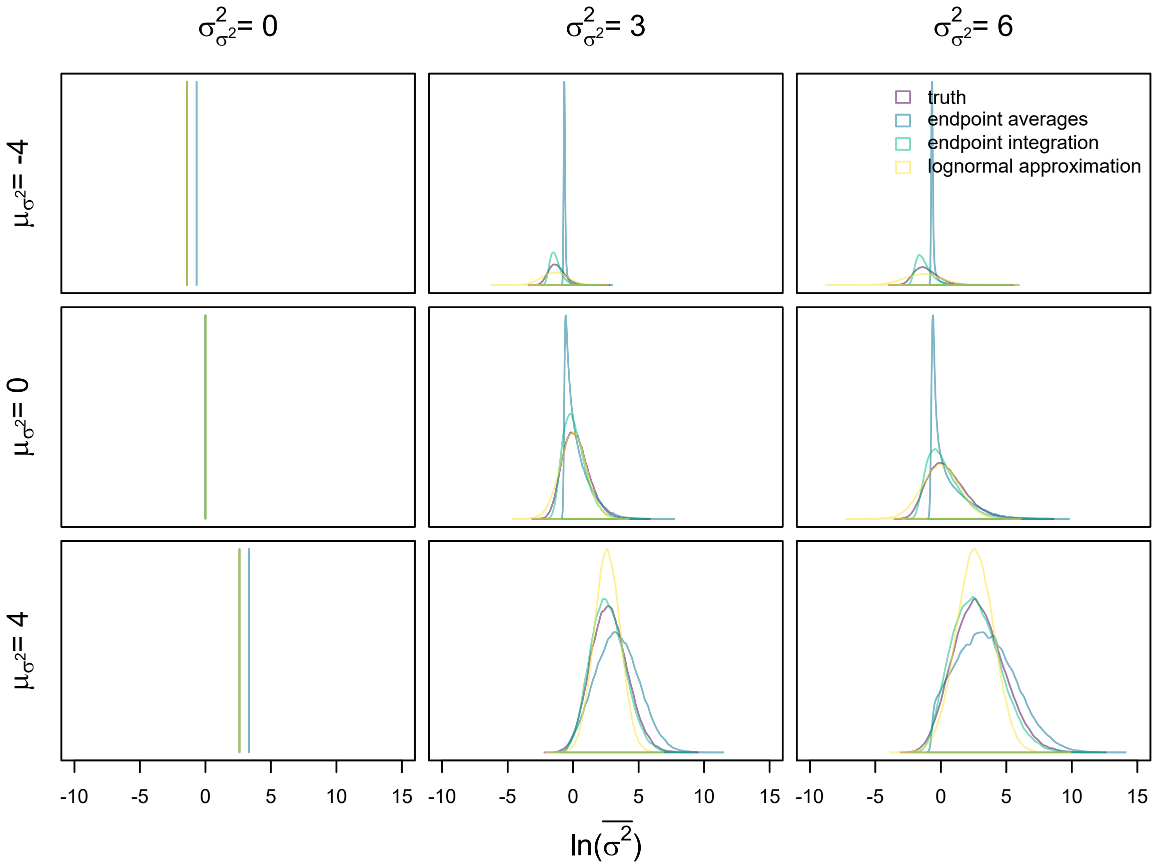


Figure S3. Distributions of simulated branchwise averages under different approximation strategies and the true distribution given parameter combinations used in the main text's simulation study. All simulations were run on single branches of length 1.

The above results help give a sense of where each approximation breaks down in parameter space, yet poorly represent the practical behavior of each approximation. In the context of our model, these approximations take place on individual branches of a phylogeny, which typically span relatively short intervals of time. For our next simulation experiment, we scaled up to simulating sets of branchwise averages on entire phylogenies. For each parameter combination (excluding combinations where rate variance is 0), we repeated the same simulations on 100 pure birth phylogenies with either 50, 100, or 200 species (generated using the R package *phytools*; Revell, 2012) standardized to a height of 1. For each phylogeny, we simulated 1,000 sets of branchwise averages under each approximation strategy, as well as fine-grained GBM sample paths ( 1,000 time points across entire phylogeny's height) representing the true distribution. Since these samples have a high number of dimensions (one for each branch in a phylogeny), we visualized how well these multivariate distributions match one another using summary statistics. Specifically, for each tree, we recorded the correlation coefficients between the means/(co)variances of branchwise averages simulated under each approximation strategy and the true distribution (Figs. S4-9). To have a null expectation for these correlation coefficients, we also simulated a second true distribution and estimated correlation coefficients for means/(co)variances between replicate true distributions.

Overall, the results indicate that all approximations do a fairly good job at recapitulating the means and (co)variances expected under the true distribution. The log-normal approximation notably exhibits uncorrelated means in the case of no trend, in contrast to other approximations. This is due to the log-normal approximation lacking the right skew of the true distribution and other approximations (Fig. 3), which naturally inflates the means of branchwise average distributions along long branches. In the case of any trend, the endpoint average approximation exhibits somewhat less strong correlations between branchwise average means compared to other approximations. When rate variance is high, the log-normal approximation exhibits performance intermediate between the endpoint average approximation and endpoint integration approximation/null distribution. However, even the worst performing simulations nearly always exhibit strong correlations in branchwise average means above 0.98. In contrast to means, correlations for branchwise average (co)variances consistently varied between about 0.98-0.99 regardless of simulation parameters or approximation strategy, closely matching the null distribution.

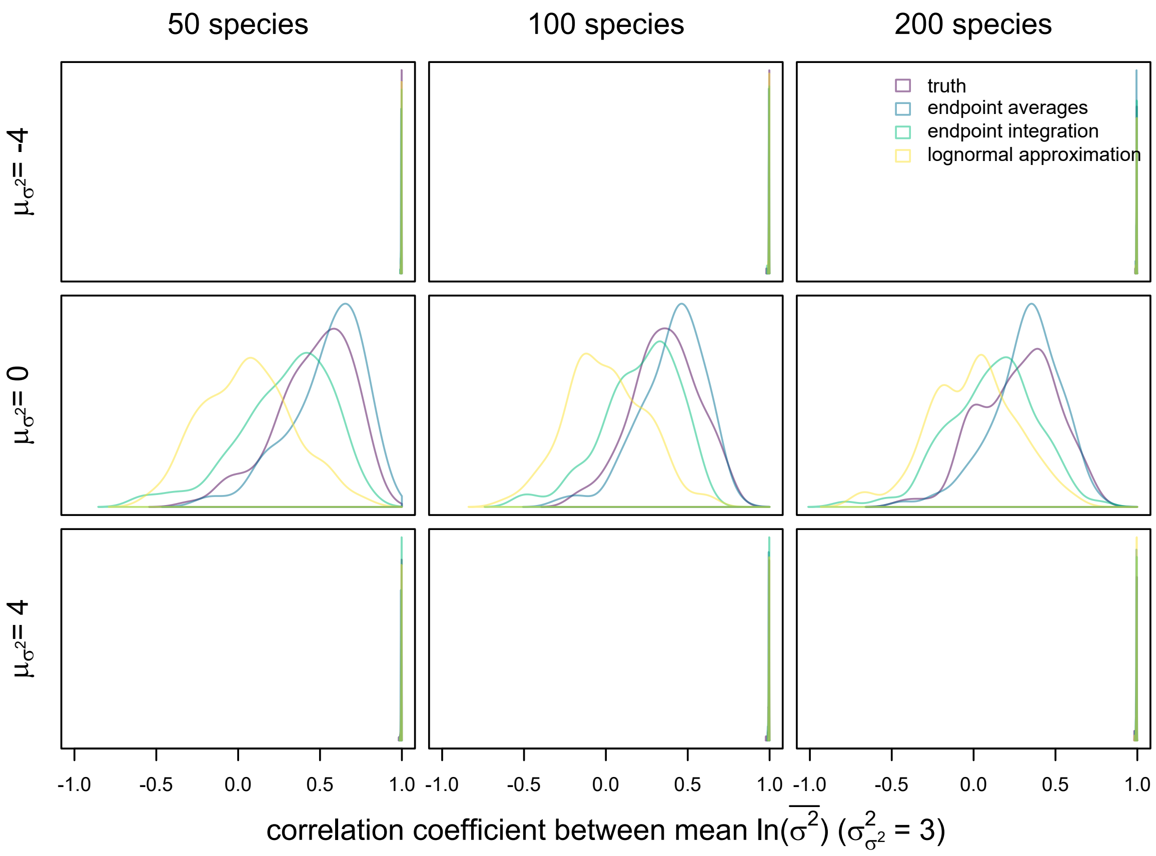


Figure S4. Distributions of correlation coefficients between mean simulated branchwise averages under different approximation strategies and the true distribution with rate variance ($\sigma^2_{\sigma^2}$) set to 3. All simulations were run on pure-birth phylogenies of height 1.

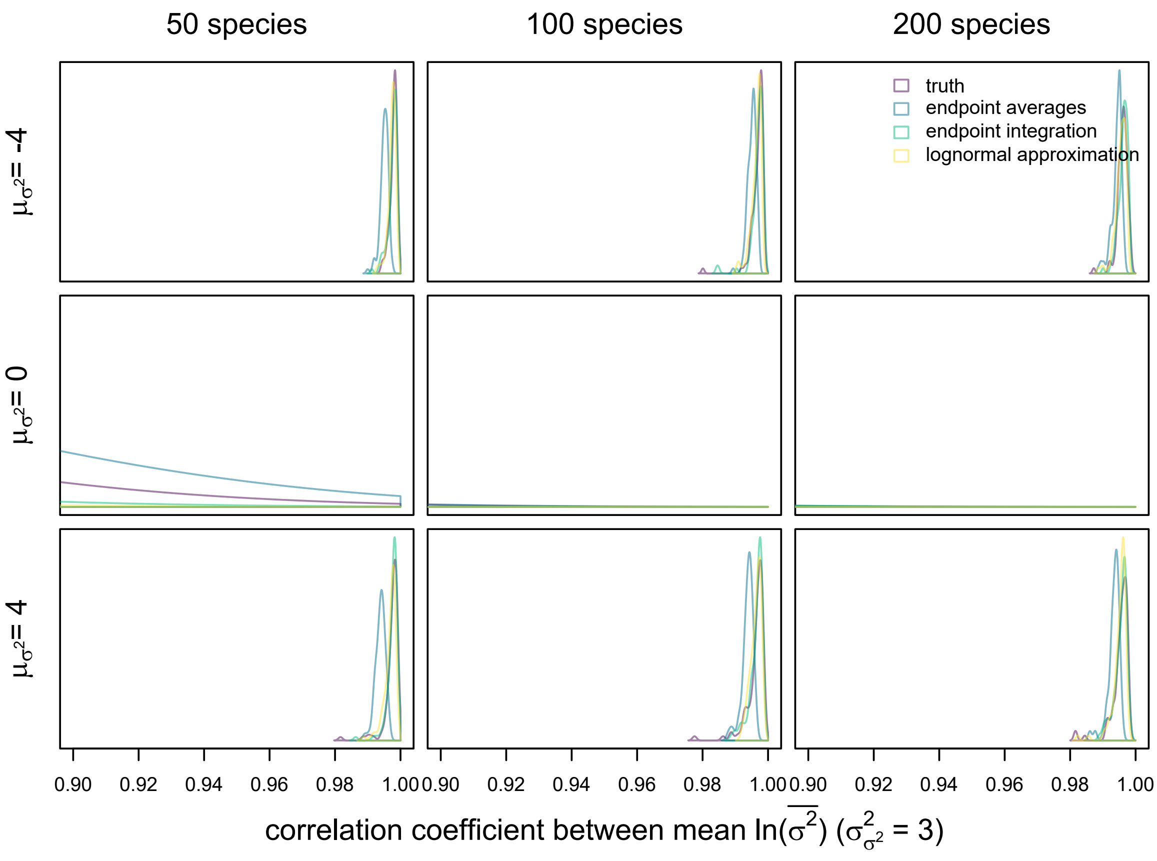


Figure S5. Distributions of correlation coefficients between mean simulated branchwise averages under different approximation strategies and the true distribution with rate variance ($\sigma^2_{\sigma^2}$) set to 3. All simulations were run on pure-birth phylogenies of height 1. Plots are zoomed in on distributions close to 1.

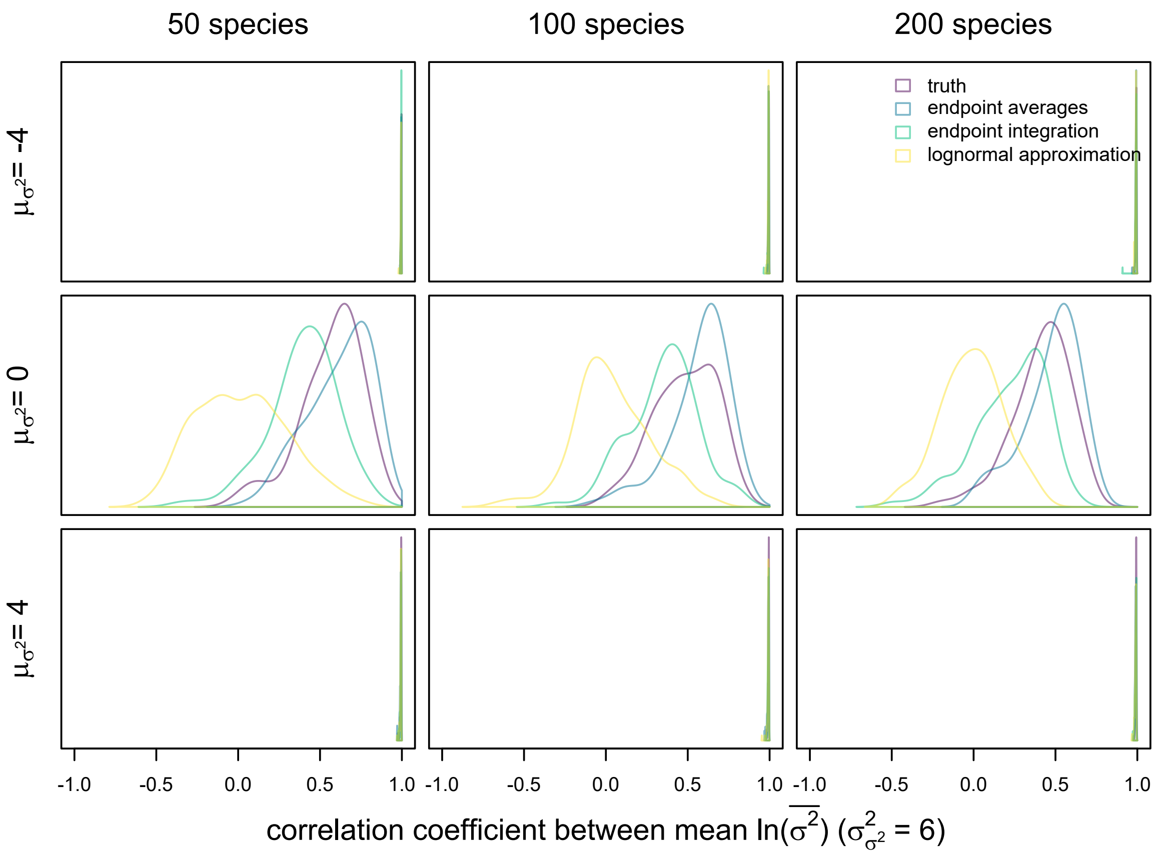


Figure S6. Distributions of correlation coefficients between mean simulated branchwise averages under different approximation strategies and the true distribution with rate variance ($\sigma^2_{\sigma^2}$) set to 6. All simulations were run on pure-birth phylogenies of height 1.

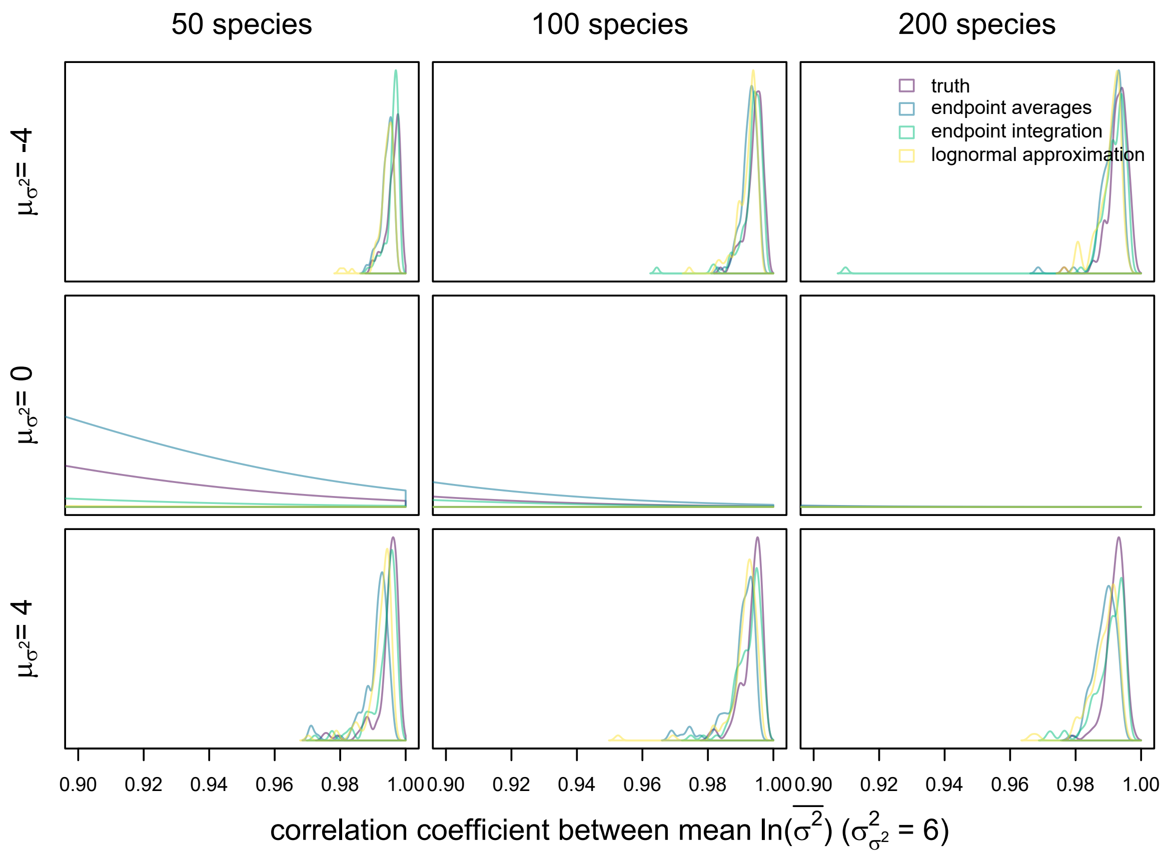


Figure S7. Distributions of correlation coefficients between mean simulated branchwise averages under different approximation strategies and the true distribution with rate variance ($\sigma^2_{\sigma^2}$) set to 6. All simulations were run on pure-birth phylogenies of height 1. Plots are zoomed in on distributions close to 1.

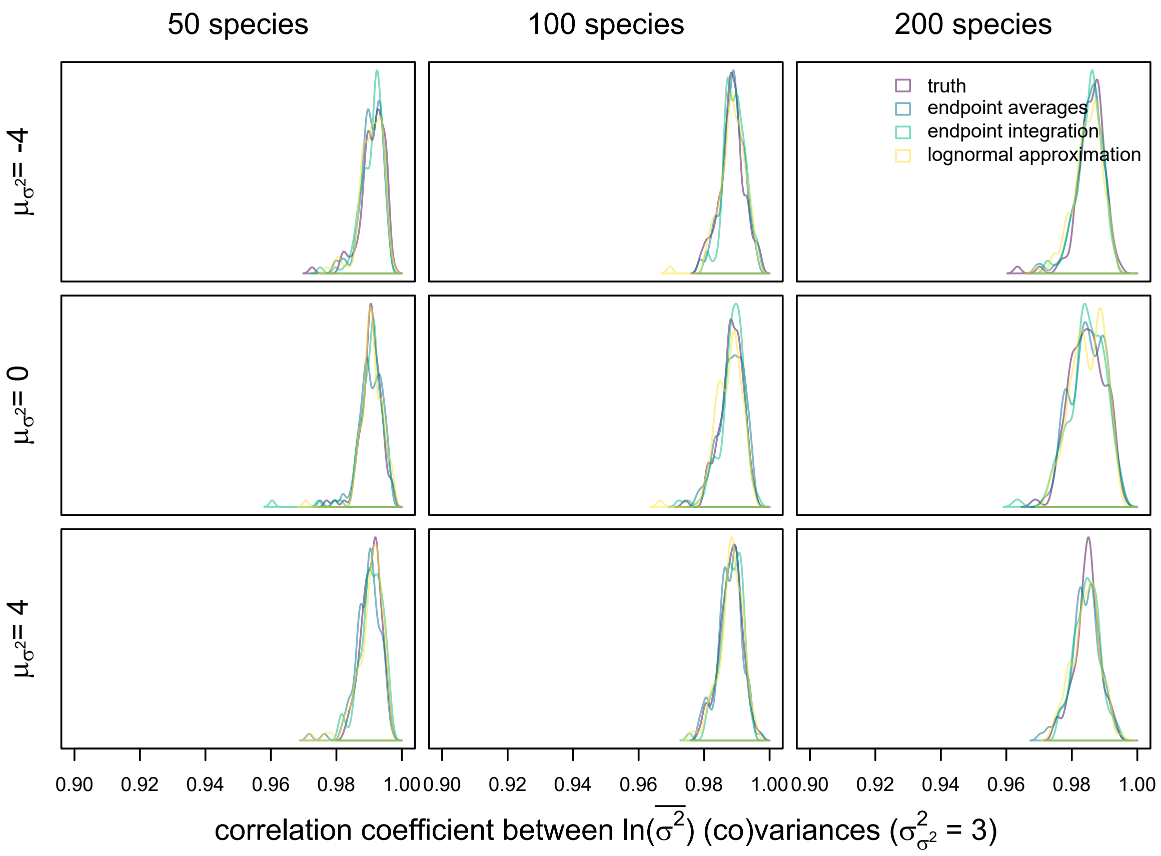


Figure S8. Distributions of correlation coefficients between simulated branchwise average (co)variances under different approximation strategies and the true distribution with rate variance ($\sigma^2_{\sigma^2}$) set to 3. All simulations were run on pure-birth phylogenies of height 1.

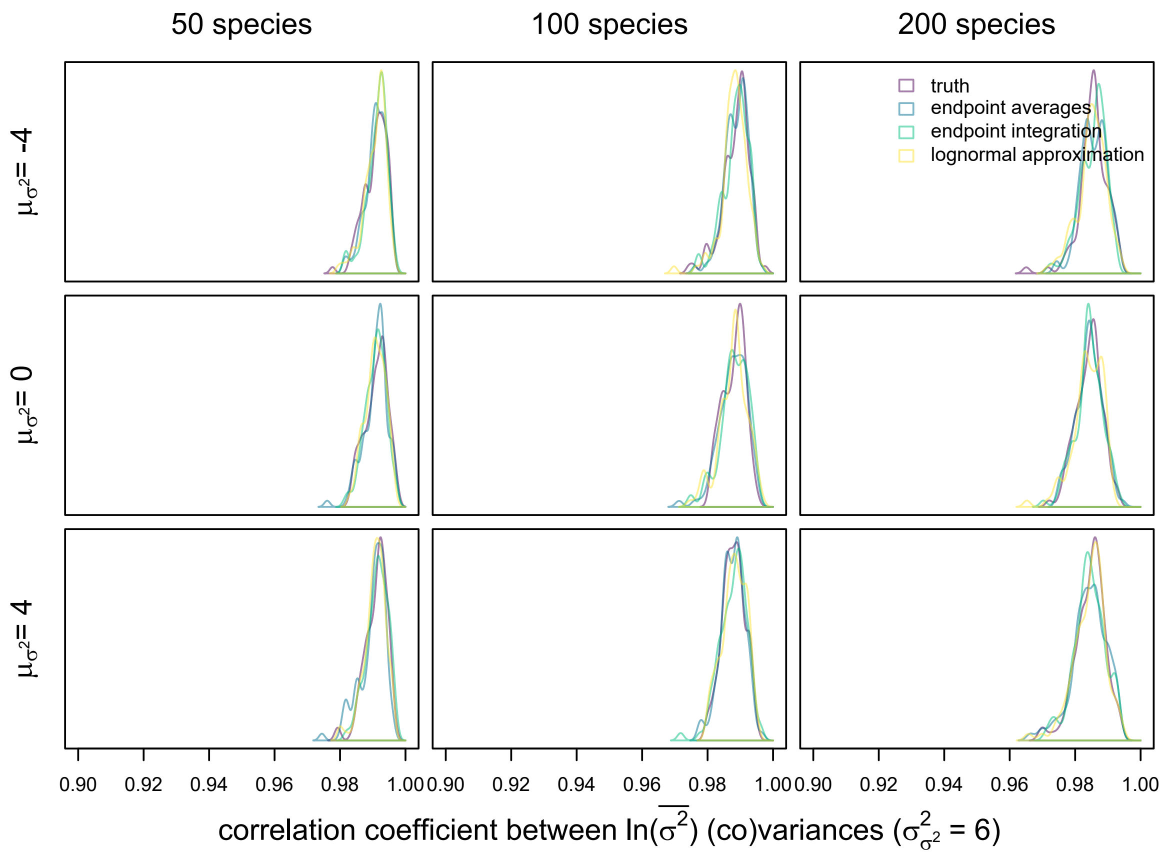


Figure S9. Distributions of correlation coefficients between simulated branchwise average (co)variances under different approximation strategies and the true distribution with rate variance ($\sigma^2_{\sigma^2}$) set to 6. All simulations were run on pure-birth phylogenies of height 1.

Since GBM time-averages are non-normally distributed, we also sought a non-parametric method of comparing samples from the approximations and true distributions. For this, we attempted to use the R package *FNN* (Beygelzimer et al., 2019) to estimate Kullback-Leibler (KL) divergence from each approximation to the true distribution. However, this estimator exhibited severe numerical issues, like negative KL divergence estimates. Thus, we instead implemented a crude K nearest neighbor probability density estimator (Zhao and Lai, 2021). For each tree in the simulation experiment above, we used this estimator to calculate local probability densities under each approximation and the true distribution around samples from a replicate true distribution. We then calculated log ratios of the true densities to densities under each approximation and averaged the distances between these log ratios and 0 (i.e., equal densities). These averaged distances give a rough sense of how well the probability density of each approximation matches that of the true distribution, with increased sampling in higher-density regions of the true distribution (Figs. S10-11). Overall, the average log density ratio distances under each approximation matches the null distribution well. The endpoint average and log-normal approximations exhibit marginally elevated distances in the case of non-zero trends and decreasing trends, respectively, likely due to these approximations' under/overestimation of branchwise average variance in certain regions of parameter space (Fig. S3).

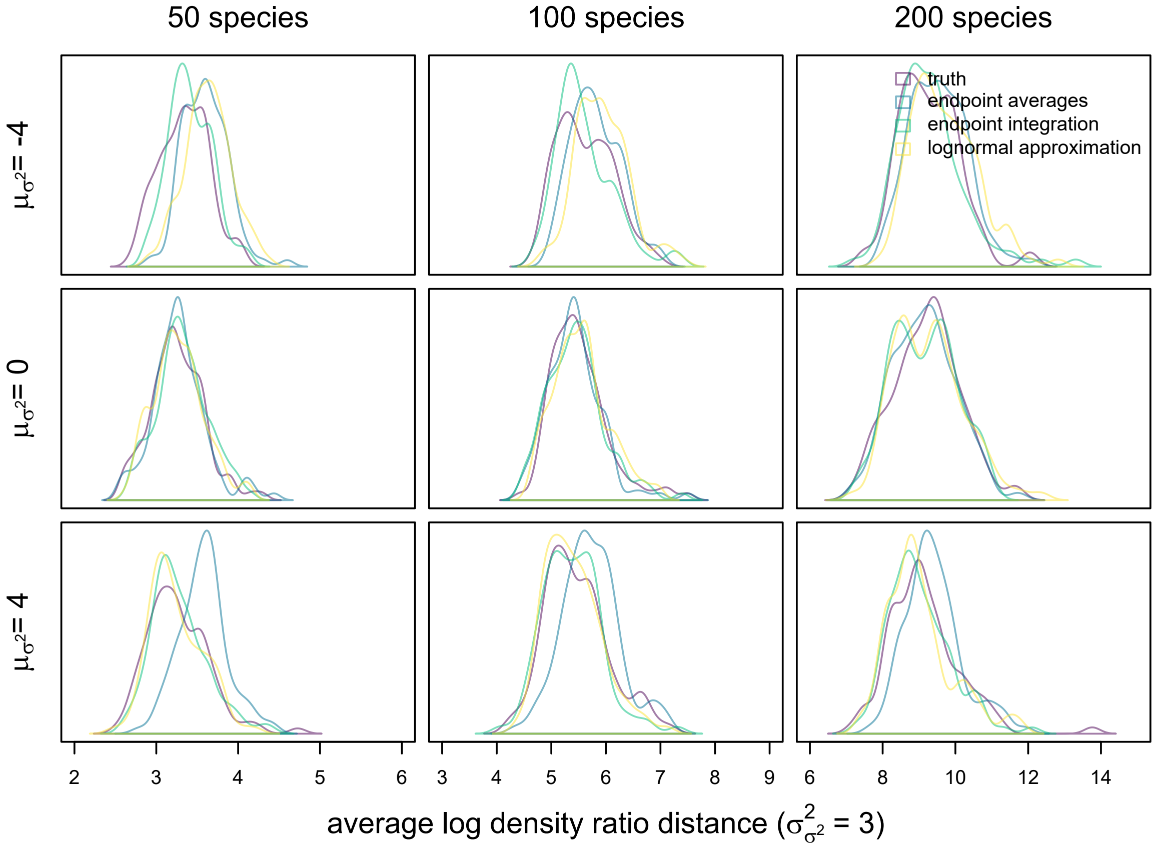


Figure S10. Distributions of average log density ratio distances between simulated branchwise average distributions under different approximation strategies and the true distribution with rate variance ($\sigma^2_{\sigma^2}$) set to 3. Probability densities were estimated via K nearest neighbors. All simulations were run on pure-birth phylogenies of height 3.

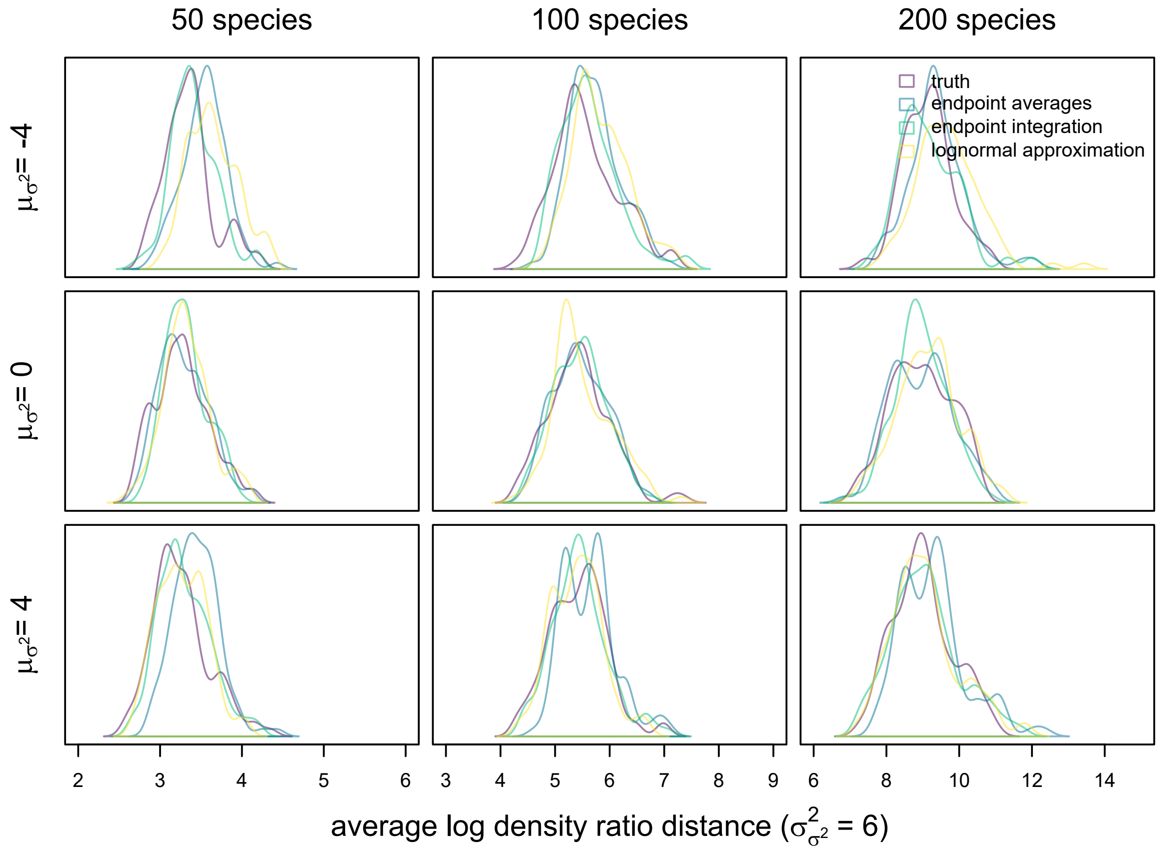


Figure S11. Distributions of average log density ratio distances between simulated branchwise average distributions under different approximation strategies and the true distribution with rate variance ($\sigma^2_{\sigma^2}$) set to 6. Probability densities were estimated via K nearest neighbors. All simulations were run on pure-birth phylogenies of height 1.

Lastly, we redid our entire simulation study with trait evolution rates simulated as evolving under a fine-grained GBM process ( 500 time points across entire phylogeny's height). We present all figures and tables for this simulation study below (Figs. S12-S16; Tables S2-4). In general, the results qualitatively match those of the simulation study presented in the main text, and we feel confident that the log-normal approximation of branchwise averages is sufficient for our model. While there is some discrepancy in the statistical power of trend detection compared to results in the main text, it is unlikely such discrepancies result from systematic bias. Notably, statistical power for trend detection even under conventional EB/LB models in this simulation study also differs from the main text results, suggesting that any discrepancies are attributable to variation in the simulated data.

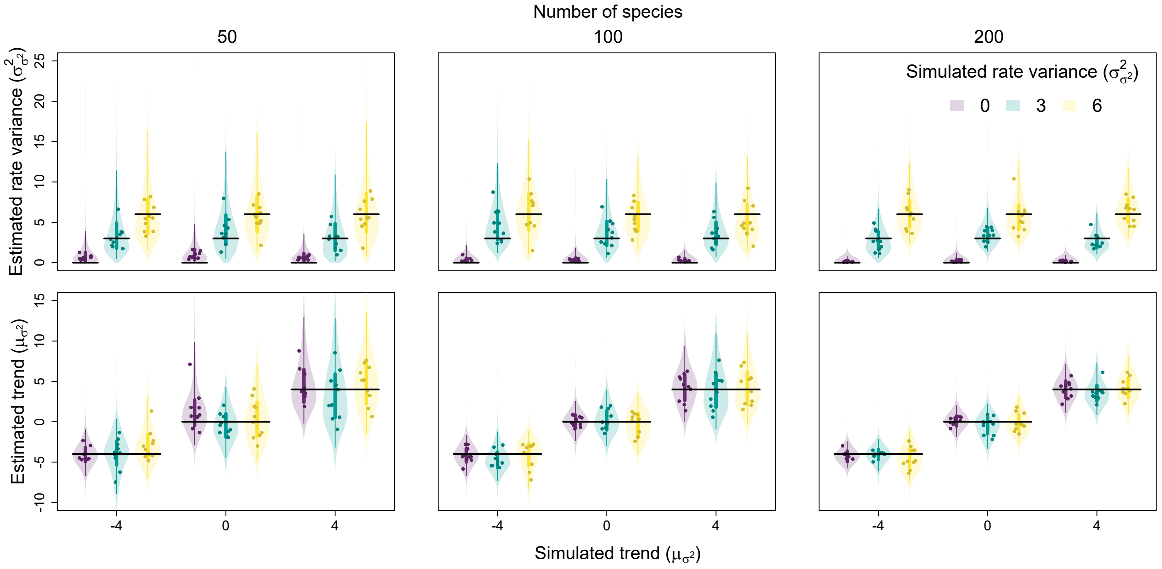


Figure S12. Relationship between simulated and estimated rate variance ($\sigma^2_{\sigma^2}$) and trend ($\mu_{\sigma^2}$) parameters. Each point is the posterior median from a single fit, while the violins are combined posterior distributions from all fits for a given trait evolution scenario. Vertical lines represent the 50% (thicker lines) and 95% credible intervals (thinner lines) of these combined posteriors, while horizontal lines represent positions of true simulated values.

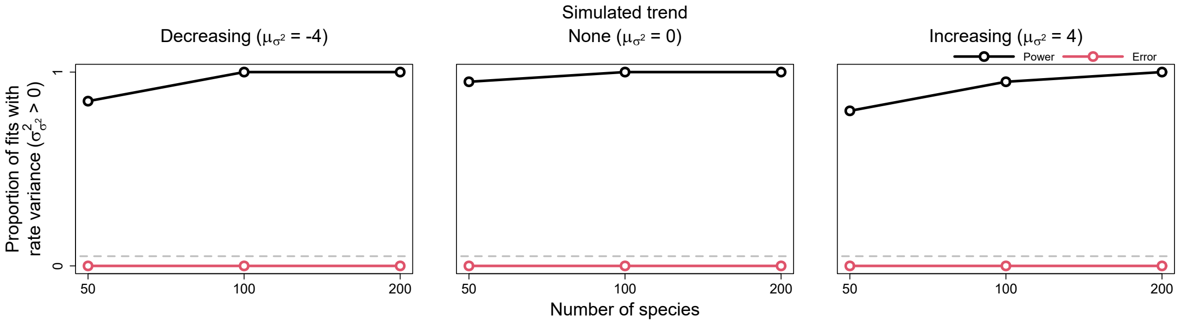


Figure S13. Power and error rates for the rate variance parameter ($\sigma^2_{\sigma^2}$). Lines depict changes in the proportion of model fits that correctly showed evidence for rate variance significantly greater than 0 (i.e., power, in black) and incorrectly showed evidence (i.e., error, in red) as a function of tree size.

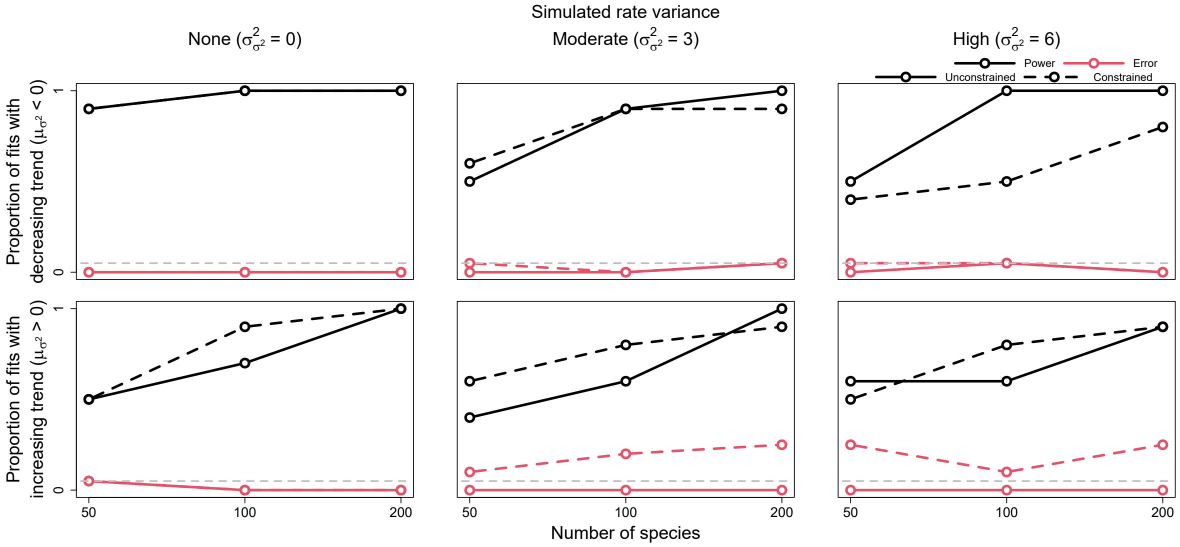


Figure S14. Power and error rates for the trend parameter ($\mu_{\sigma^2}$). Lines depict changes in the proportion of model fits that correctly showed evidence for trends significantly less and greater than 0 (i.e., power, in black) and incorrectly showed evidence (i.e., error, in red) as a function of tree size. Results are shown for both models allowed to freely estimate rate variance ($\sigma^2_{\sigma^2}$) (i.e., unconstrained models, solid lines) and models with rate variance constrained to 0 (i.e., constrained models, dashed lines). The latter models are identical to conventional early/late burst models.

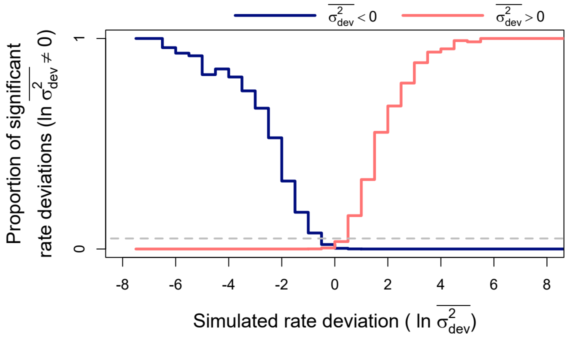


Figure S15. Power and error rates for branchwise rate parameters (ln $\overline{\sigma^2}$). Lines depict changes in proportions of branchwise rates considered anomalously slow (in blue) or fast (in red) as a function of simulated rate deviations (ln $\overline{\sigma^2_{dev}}$). These results combine all fits to simulated data that detected rate variance ($\sigma^2_{\sigma^2}$) significantly greater than 0. The proportions are equivalent to power when the detected rate deviation is of the same sign as the true, simulated deviation (left of 0 for anomalously slow rates in blue and right for anomalously fast rates in red), and to error rate when the detected and true rate deviations are of opposite signs. Here, significant rate deviations for simulated rate deviations that are exactly 0 are considered errors regardless of sign.

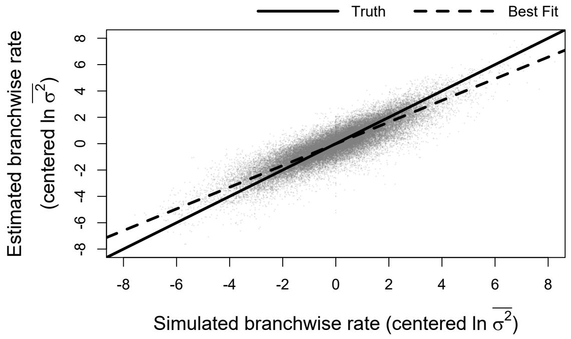


Figure S16. Relationship between simulated and estimated branchwise rate parameters (ln $\overline{\sigma^2}$). For each simulation and posterior sample, branchwise rates were first centered by subtracting their mean. We estimated centered branchwise rates by taking the median of the centered posterior samples. The solid line represents the position of the true centered branchwise rates, while the shallower, dashed line represents the observed line of best fit for these data.

Table S2. *Average posterior relative accuracy for rate variance, trend, and branchwise rate parameters for each simulated trait evolution scenario and tree size*

| | | rate variance | | | trend | | | branchwise rates | | |
|---|---|---|---|---|---|---|---|---|---|---|
| $\sigma^2_{\sigma^2}$ = | | 0 | 3 | 6 | 0 | 3 | 6 | 0 | 3 | 6 |
| | | | | | | 50 species | | | | |
| $\mu_{\sigma^2}$ = | -4 | 0.16 | 0.10 | 0.12 | 0.12 | 0.21 | 0.18 | 0.11 | 0.19 | 0.21 |
| | 0 | 0.20 | 0.13 | 0.12 | 0.22 | 0.17 | 0.24 | 0.17 | 0.18 | 0.21 |
| | 4 | 0.17 | 0.14 | 0.13 | 0.13 | 0.26 | 0.17 | 0.16 | 0.22 | 0.19 |
| | | | | | | 100 species | | | | |
| $\mu_{\sigma^2}$ = | -4 | 0.16 | 0.20 | 0.28 | 0.21 | 0.24 | 0.27 | 0.12 | 0.18 | 0.19 |
| | 0 | 0.18 | 0.20 | 0.16 | 0.14 | 0.19 | 0.22 | 0.11 | 0.21 | 0.22 |
| | 4 | 0.18 | 0.17 | 0.29 | 0.21 | 0.25 | 0.23 | 0.16 | 0.21 | 0.21 |
| | | | | | | 200 species | | | | |
| $\mu_{\sigma^2}$ = | -4 | 0.20 | 0.31 | 0.20 | 0.16 | 0.06 | 0.30 | 0.12 | 0.22 | 0.20 |
| | 0 | 0.19 | 0.18 | 0.24 | 0.11 | 0.21 | 0.23 | 0.09 | 0.20 | 0.21 |
| | 4 | 0.21 | 0.24 | 0.17 | 0.19 | 0.18 | 0.13 | 0.13 | 0.22 | 0.19 |

Table S3. *Average posterior breadth for rate variance, trend, and branchwise rate parameters for each simulated trait evolution scenario and tree size*

| | | rate variance | | | trend | | | branchwise rates | | |
|---|---|---|---|---|---|---|---|---|---|---|
| $\sigma^2_{\sigma^2} =$ | | 0 | 3 | 6 | 0 | 3 | 6 | 0 | 3 | 6 |
| | | | | | 50 species | | | | | |
| $\mu_{\sigma^2} =$ | -4 | 3.67 | 9.11 | 12.98 | 4.66 | 6.02 | 6.81 | 2.28 | 3.24 | 3.65 |
| | 0 | 4.38 | 10.67 | 12.60 | 7.28 | 7.09 | 8.00 | 2.60 | 3.41 | 3.89 |
| | 4 | 3.35 | 9.00 | 13.88 | 10.34 | 10.95 | 12.09 | 2.81 | 3.50 | 4.10 |
| | | | | | 100 species | | | | | |
| $\mu_{\sigma^2} =$ | -4 | 1.77 | 7.96 | 9.58 | 3.53 | 4.56 | 4.72 | 1.71 | 3.22 | 3.46 |
| | 0 | 1.64 | 6.72 | 9.15 | 4.04 | 5.09 | 5.67 | 1.76 | 3.12 | 3.42 |
| | 4 | 1.36 | 6.77 | 8.13 | 6.74 | 8.08 | 7.86 | 1.87 | 3.31 | 3.55 |
| | | | | | 200 species | | | | | |
| $\mu_{\sigma^2} =$ | -4 | 0.71 | 3.97 | 7.20 | 2.64 | 3.58 | 4.06 | 1.24 | 2.50 | 3.12 |
| | 0 | 1.04 | 4.26 | 6.52 | 3.34 | 3.98 | 4.15 | 1.36 | 2.77 | 3.25 |
| | 4 | 0.79 | 3.62 | 6.89 | 4.53 | 4.88 | 5.69 | 1.39 | 2.70 | 3.37 |

Table S4. *Average posterior coverage for rate variance, trend, and branchwise rate parameters for each simulated trait evolution scenario and tree size*

| | | rate variance | | | trend | | | branchwise rates | | |
|---|---|---|---|---|---|---|---|---|---|---|
| $\sigma^2_{\sigma^2} =$ | | 0 | 3 | 6 | 0 | 3 | 6 | 0 | 3 | 6 |
| | | | | | 50 species | | | | | |
| $\mu_{\sigma^2} =$ | -4 | — | 1.00 | 1.00 | 1.00 | 0.80 | 0.90 | 1.00 | 0.95 | 0.94 |
| | 0 | — | 1.00 | 1.00 | 0.90 | 1.00 | 1.00 | 0.97 | 0.98 | 0.94 |
| | 4 | — | 1.00 | 1.00 | 1.00 | 0.80 | 1.00 | 0.95 | 0.94 | 0.96 |
| | | | | | 100 species | | | | | |
| $\mu_{\sigma^2} =$ | -4 | — | 0.70 | 0.90 | 0.90 | 1.00 | 0.90 | 1.00 | 0.96 | 0.96 |
| | 0 | — | 1.00 | 1.00 | 1.00 | 1.00 | 0.90 | 1.00 | 0.94 | 0.94 |
| | 4 | — | 1.00 | 0.90 | 0.90 | 0.90 | 1.00 | 0.99 | 0.95 | 0.93 |
| | | | | | 200 species | | | | | |
| $\mu_{\sigma^2} =$ | -4 | — | 0.90 | 1.00 | 1.00 | 1.00 | 0.90 | 0.99 | 0.93 | 0.95 |
| | 0 | — | 1.00 | 0.80 | 1.00 | 0.90 | 1.00 | 1.00 | 0.95 | 0.95 |
| | 4 | — | 1.00 | 1.00 | 0.90 | 1.00 | 1.00 | 0.99 | 0.93 | 0.96 |

### Average Changes in Trait Evolution Rates

Conventional EB/LB models of trait evolution assume that rates follow a homogeneous, exponential declines or increases with respect to time (Blomberg et al., 2003). The definition of EBs/LBs under such models is thus straight-forward–any given time slice in a clade's history is associated with a single trait evolution rate, and these rates can only decrease, increase or stay the same. On the other hand, allowing for rate heterogeneity independent of overall temporal trends means that any given time slice in a clade's history is associated with a *distribution* of trait evolution rates. Because of this, our new method allows for alternative definitions of EBs/LBs, depending on how one summarizes these distributions. In the current study, we mainly consider a definition based on whether the medians, or geometric means, of these distributions decrease or increase over time (change per unit time given by $\mu_{\sigma^2}$, hereafter the "trend" parameter, as in the main text). Alternatively, one could use a definition based on whether the average, or arithmetic means, of these distributions decrease or increase over time (change per unit time given by $\mu_{\sigma^2} + \sigma^2_{\sigma^2}/2$, hereafter the "average change" parameter, $\delta_{\sigma^2}$).

We chose to focus on trend over average change estimation and define EBs/LBs based on the trend parameter for a few reasons. First, average change is a composite parameter of both the trend and rate variance parameters, posing some interpretational challenges. In general, it seems more intuitive to consider the magnitude of deterministic changes in trait evolution rates (the trend component) apart from the magnitude of stochastic changes (the rate variance component). Second, since rates evolve in an approximately log-normal manner under our model, medians are a natural, reliable way of summarizing their distributions, corresponding to the exponentiated average of rates on the natural log scale. In contrast, the right skew of log-normal distributions causes raw averages of trait evolution rates to be highly influenced by few, extreme outliers, particularly when rate variance is high. For this reason, our model can produce trait evolution scenarios whereby rates exhibit declines in the majority of lineages (directly related to changes in median rates) while increasing on average (Figs. S17-18). Lastly, many macroevolutionary biologists consider “accounting” for lineages/subclades exhibiting unusual trait evolution rates critical to elucidating and understanding changes in rates over time (Lloyd et al., 2012; Slater and Pennell, 2014; Benson et al., 2014; Hopkins and Smith, 2015; Wright, 2017; Puttick, 2018). This implies that many empiricists intuitively define EBs/LBs based on majority changes in rates rather than changes in average rates. Additionally, by log-transforming traits prior to analysis, many macroevolutionary biologists implicitly use GBM processes to model trait evolution, just as we use a (approximate) GBM process to model rate evolution here. In the context of trait evolution, the analogous trend parameter is widely considered by empiricists and method developers alike to determine whether a clade exhibits a directional “evolutionary trend” in traits, regardless the estimated variance parameter (Hunt, 2006; Raj Pant et al., 2014; Sookias et al., 2012; Gill et al., 2017).

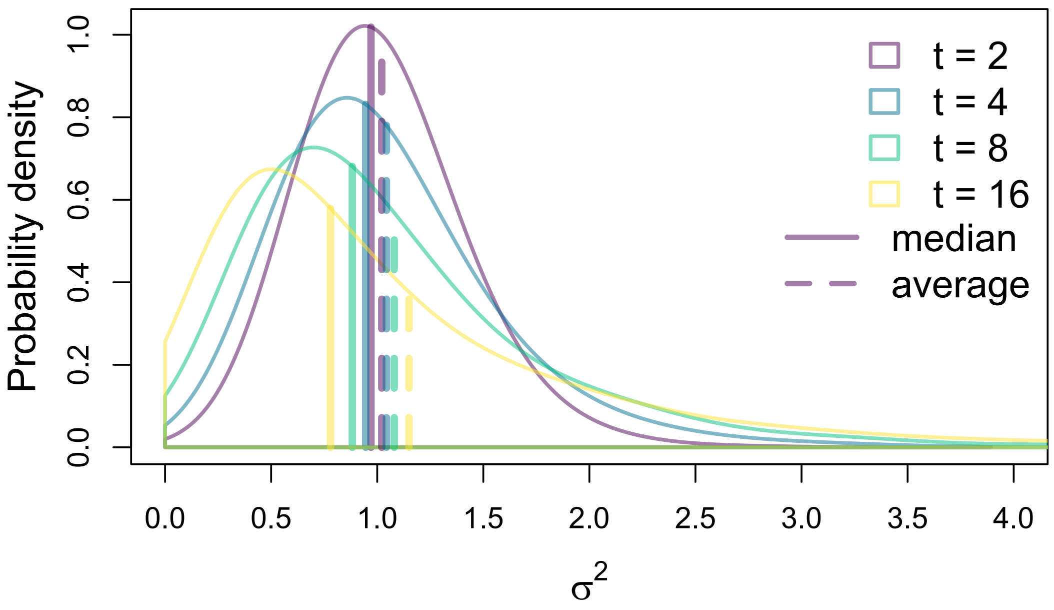


Figure S17. Distributions of 6,000 rates simulated as evolving under a GBM process with trend of -0.015 and rate variance of 0.05 at various time points, with starting rate of 1 at time $t = 0$. Parameter values were chosen to clearly illustrate how rates under our model may exhibit majority declines while increasing on average due to the skewed nature of rate change. Solid and dashed vertical lines represent the positions of median and average rate values, respectively, for each time point.

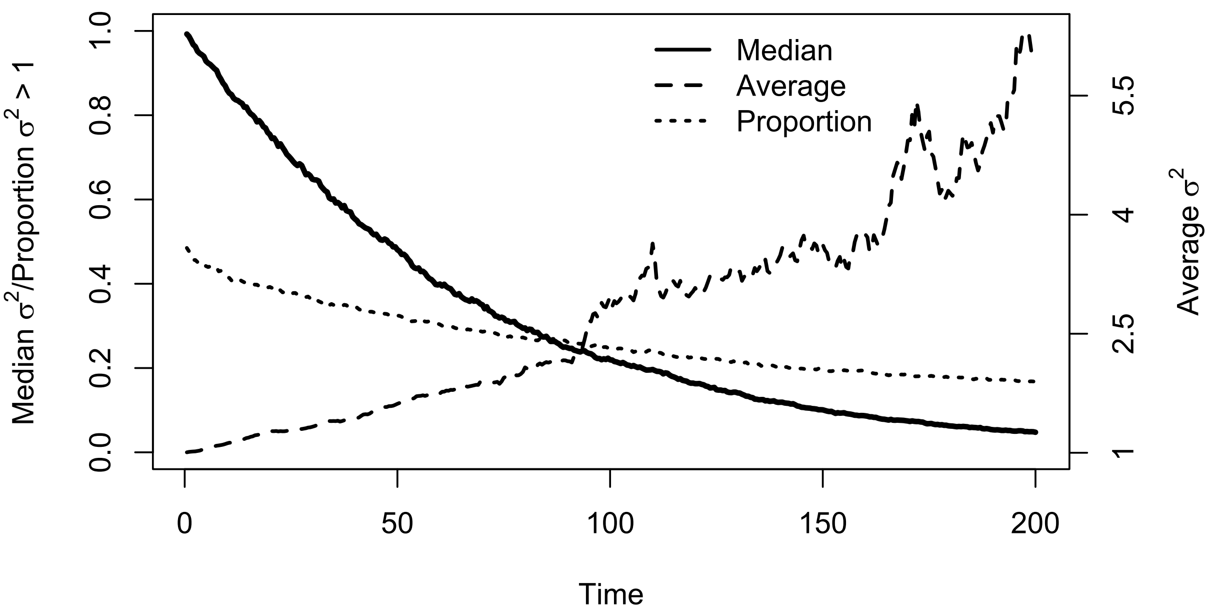


Figure S18. Changes over time in the median and average of 6,000 rates simulated as evolving under a GBM process with trend of -0.015 and rate variance of 0.05, with starting rate of 1 at time $t = 0$. Parameter values were chosen to clearly illustrate how rates under our model may exhibit majority declines while increasing on average due to the skewed nature of rate change. Solid and dashed lines depict changes in median and average rate values, respectively, while the dotted line depicts changes in the proportion of rates less than the starting rate of 1.

Here, we briefly consider our new method's performance with respect to estimating and detecting average changes in trait evolution rates. Interestingly, our simulation study results revealed that, in the presence of time-independent rate heterogeneity, conventional EB/LB models (equivalent to our new models with rate variance constrained to 0) appear to estimate average change, rather than trend parameters, as defined under our model (Figs. S19-20). We are not aware of any previous research explicitly demonstrating this phenomenon. When comparing performance of constrained to unconstrained models with respect to detecting significant average change (i.e., 95% credible interval lies entirely below or above 0), we see only a modest reduction in error rates and greatly reduced power to detect negative average change under the full, unconstrained model (Fig. S21). Nonetheless, inference of the average change parameter seems substantially improved under unconstrained models (Tables S5-7). In the presence of time-independent rate heterogeneity, constrained models exhibit less accurate, overly-narrow posterior estimates of average change, resulting in rather low posterior coverage. This warrants caution in interpreting the results of conventional EB/LB models fitted to comparative data exhibiting substantial time-independent rate heterogeneity, and we recommend estimating rate variance even when one's only goal is to estimate changes in average trait evolution rates over time.

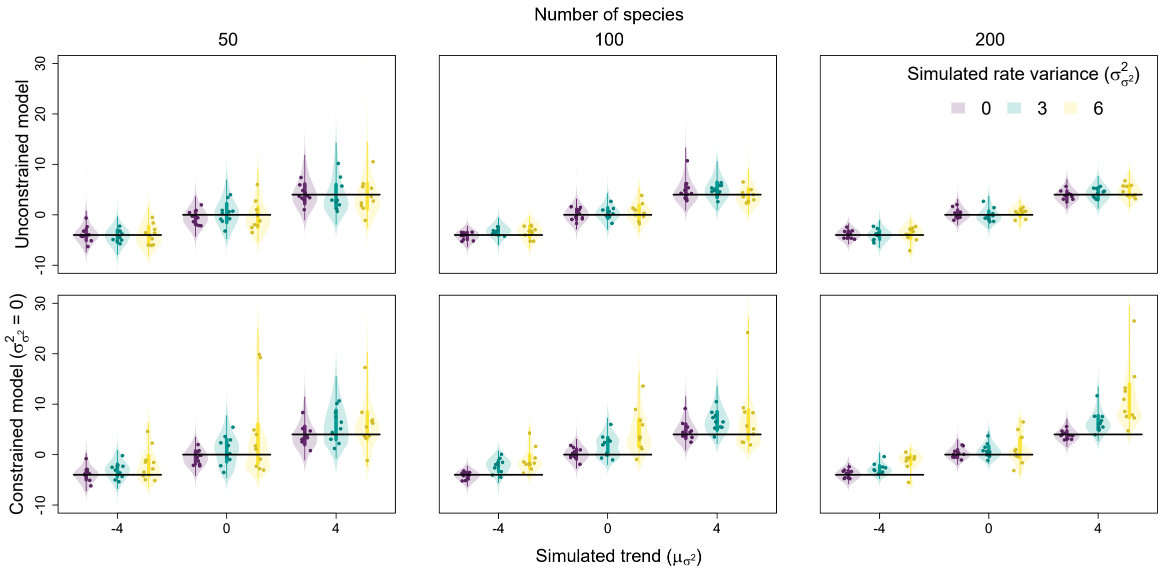


Figure S19. Relationship between simulated rate variance ($\sigma^2_{\sigma^2}$)/trend ($\mu_{\sigma^2}$) and estimated trend parameters. Each point is the posterior median from a single fit, while the violins are combined posterior distributions from all fits for a given trait evolution scenario. Vertical lines represent the 50% (thicker lines) and 95% credible intervals (thinner lines) of these combined posteriors, while horizontal lines represent positions of true simulated values. Results for models with estimated rate variance unconstrained and constrained to 0 are shown on top and bottom, respectively.

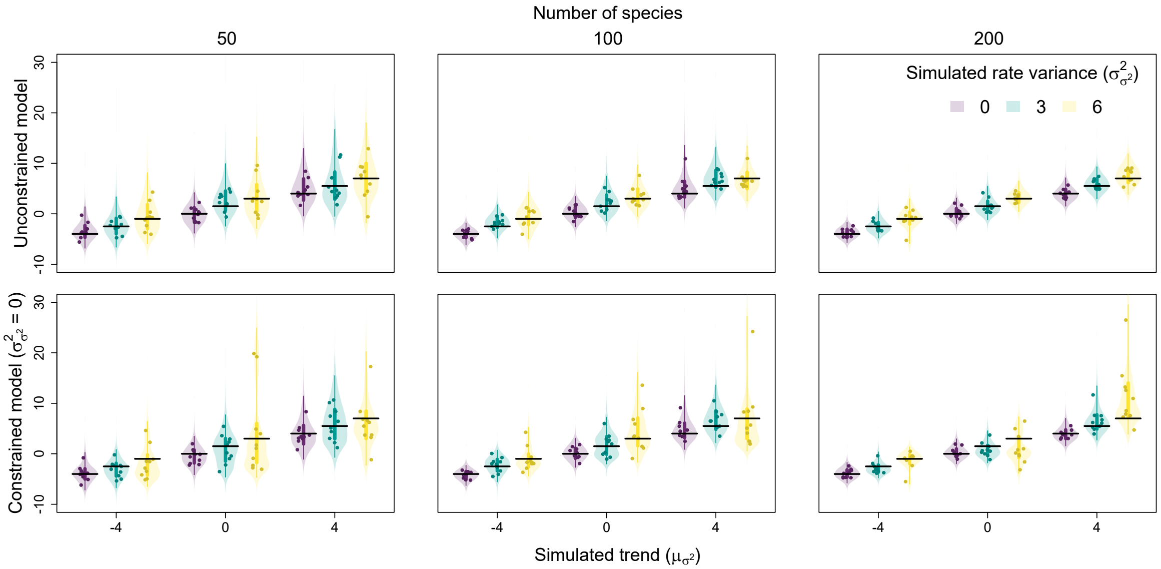


Figure S20. Relationship between simulated rate variance ($\sigma^2_{\sigma^2}$)/trend ($\mu_{\sigma^2}$) and estimated average change ($\delta_{\sigma^2}$) parameters. Each point is the posterior median from a single fit, while the violins are combined posterior distributions from all fits for a given trait evolution scenario. Vertical lines represent the 50% (thicker lines) and 95% credible intervals (thinner lines) of these combined posteriors, while horizontal lines represent positions of true simulated values. Results for models with estimated rate variance unconstrained and constrained to 0 are shown on top and bottom, respectively.

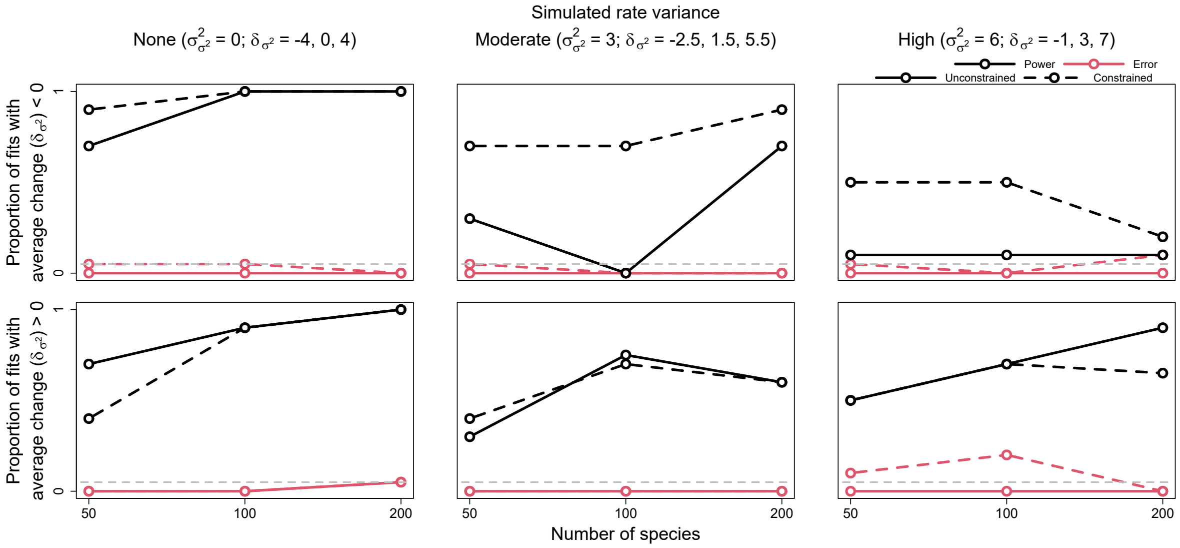


Figure S21. Power and error rates for the average parameter ($\delta_{\sigma^2}$). Lines depict changes in the proportion of model fits that correctly showed evidence for average change significantly less and greater than 0 (i.e., power, in black) and incorrectly showed evidence (i.e., error, in red) as a function of tree size. Results are shown for both models allowed to freely estimate rate variance ($\sigma^2_{\sigma^2}$) (i.e., unconstrained models, solid lines) and models with rate variance constrained to 0 (i.e., constrained models, dashed lines). The latter models are identical to conventional early/late burst models.

Table S5. *Average posterior relative accuracy for average change parameter under models with rate variance unconstrained and constrained to 0 for each simulated trait evolution scenario and tree size*

| | | unconstrained | | | constrained | | |
|---|---|---|---|---|---|---|---|
| $\sigma^2_{\sigma^2} =$ | | 0 | 3 | 6 | 0 | 3 | 6 |
| | | 50 species | | | | | |
| $\mu_{\sigma^2} =$ | -4 | 0.17 | 0.14 | 0.18 | 0.19 | 0.27 | 0.49 |
| | 0 | 0.18 | 0.17 | 0.22 | 0.22 | 0.37 | 0.68 |
| | 4 | 0.12 | 0.20 | 0.26 | 0.15 | 0.26 | 0.34 |
| | | 100 species | | | | | |
| $\mu_{\sigma^2} =$ | -4 | 0.16 | 0.17 | 0.23 | 0.18 | 0.35 | 0.44 |
| | 0 | 0.21 | 0.21 | 0.15 | 0.23 | 0.41 | 0.43 |
| | 4 | 0.17 | 0.16 | 0.13 | 0.16 | 0.19 | 0.53 |
| | | 200 species | | | | | |
| $\mu_{\sigma^2} =$ | -4 | 0.25 | 0.17 | 0.23 | 0.26 | 0.30 | 0.34 |
| | 0 | 0.20 | 0.20 | 0.12 | 0.21 | 0.41 | 0.91 |
| | 4 | 0.13 | 0.14 | 0.19 | 0.14 | 0.24 | 0.44 |

Table S6. *Average posterior breadth for average change parameter under models with rate variance unconstrained and constrained to 0 for each simulated trait evolution scenario and tree size*

| | | unconstrained | | | constrained | | |
|---|---|---|---|---|---|---|---|
| $\sigma^2_{\sigma^2} =$ | | 0 | 3 | 6 | 0 | 3 | 6 |
| | | 50 species | | | | | |
| $\mu_{\sigma^2} =$ | -4 | 5.56 | 7.95 | 10.27 | 4.50 | 4.65 | 5.63 |
| | 0 | 6.25 | 9.89 | 11.46 | 5.47 | 6.82 | 8.45 |
| | 4 | 11.04 | 11.69 | 12.45 | 9.48 | 10.81 | 10.60 |
| | | 100 species | | | | | |
| $\mu_{\sigma^2} =$ | -4 | 3.42 | 5.48 | 6.54 | 3.07 | 3.49 | 3.84 |
| | 0 | 4.46 | 6.27 | 7.44 | 3.97 | 4.58 | 6.60 |
| | 4 | 7.64 | 8.97 | 8.56 | 7.12 | 8.41 | 8.40 |
| | | 200 species | | | | | |
| $\mu_{\sigma^2} =$ | -4 | 2.82 | 4.14 | 5.07 | 2.70 | 2.87 | 3.26 |
| | 0 | 3.40 | 4.50 | 5.13 | 3.25 | 3.30 | 3.37 |
| | 4 | 4.51 | 5.45 | 6.29 | 4.38 | 5.78 | 8.73 |

Table S7. *Average posterior coverage for average change parameter under models with rate variance unconstrained and constrained to 0 for each simulated trait evolution scenario and tree size*

| | | unconstrained | | | constrained | | |
|---|---|---|---|---|---|---|---|
| $\sigma^2_{\sigma^2} =$ | | 0 | 3 | 6 | 0 | 3 | 6 |
| | | 50 species | | | | | |
| $\mu_{\sigma^2} =$ | -4 | 0.90 | 1.00 | 0.90 | 0.80 | 0.60 | 0.50 |
| | 0 | 1.00 | 1.00 | 1.00 | 0.90 | 0.80 | 0.40 |
| | 4 | 1.00 | 1.00 | 0.90 | 1.00 | 1.00 | 0.80 |
| | | 100 species | | | | | |
| $\mu_{\sigma^2} =$ | -4 | 1.00 | 1.00 | 0.90 | 1.00 | 0.80 | 0.60 |
| | 0 | 1.00 | 0.90 | 0.90 | 0.90 | 0.60 | 0.60 |
| | 4 | 0.90 | 1.00 | 1.00 | 0.90 | 0.90 | 0.60 |
| | | 200 species | | | | | |
| $\mu_{\sigma^2} =$ | -4 | 1.00 | 1.00 | 0.90 | 1.00 | 0.90 | 0.80 |
| | 0 | 0.90 | 1.00 | 1.00 | 0.90 | 0.60 | 0.10 |
| | 4 | 1.00 | 1.00 | 1.00 | 1.00 | 0.90 | 0.50 |
